# Regulation of Crassulacean Acid Metabolism at the protein level in the CAM plant *Kalanchoë laxiflora*

**DOI:** 10.1101/2024.10.25.620298

**Authors:** Katharina Schiller, Saskia Janshoff, Sanja Zenker, Prisca Viehöver, James Hartwell, Jürgen Eirich, Iris Finkemeier, Andrea Bräutigam

## Abstract

Crassulacean acid metabolism (CAM) is an adaptation to environments where water availability is seasonal or extremely low. It serves to ensure survival and/ or maintain productivity in these adverse environments. CAM has repeatedly evolved although it requires a large and complex set of enzymes and transporters and regulatory processes to control metabolite flux and pools. To test potential regulatory levels, we analyze the CAM plant *Kalanchoë laxiflora* embedded in the context of available CAM genome and transcriptome sequences. We show that CAM associated transcripts and proteins do not show a binary on/off pattern between day and night in *K. laxiflora*. Instead, we observe that many CAM plants display shared amino acid changes compared to C_3_ plants especially in starch metabolism. Phosphoproteomics identifies phosphoproteome changes in *K. laxiflora* between day and night. Taken together, the analyses demonstrate the CAM photosynthesis is regulated at the levels of transcripts and proteins.

**One sentence summary:** Regulation of CAM cannot be explained by transcript and protein abundance alone but is also dependent on adaptive changes in proteins and posttranslational modifications.

## Introduction

Crassulacean acid metabolism (CAM) is an adaptation to dry or seasonally dry environments and ensures survival and/ or maintains productivity in these adverse environments. CAM has evolved independently and repeatedly in many lineages of plants, even though it requires a large and complex set of enzymes and transporters as well as regulatory processes to control metabolite flux and pools (Schiller and Bräutigam, 2021). CAM plants separate primary uptake and fixation of atmospheric carbon dioxide (CO_2_) in the dark from subsequent secondary re-fixation in the Calvin Benson Bassham (CBB) cycle in the light. They take advantage of cooler temperatures and higher relative humidity at night, which allows them to lose less water through their open stomata during primary atmospheric CO_2_ assimilation (Dodd et al., 2002; Borland et al., 2014). CAM plants use the enzyme phospho*enol*pyruvate (PEP) carboxylase (PPC) for nocturnal CO_2_ assimilation. PPC converts PEP plus HCO_3_^-^ to 4-carbon oxaloacetate (OAA), and OAA is rapidly reduced to malate by malate dehydrogenase (MDH). Malate is stored as malic acid in the vacuole. During sunrise, malic acid begins to be released from the vacuole, and is decarboxylated by either NAD(P)-malic enzyme (NAD(P)-ME), or MDH combined with PEP carboxykinase (PEP-CK). The specific decarboxylation pathway used depends on the species. Malate decarboxylation in the light releases CO_2_ for use by ribulose 1,5-bisphosphate carboxylase oxygenase (Rubisco), the first step of the CBB cycle (Ludwig et al., 2024). The remaining three-carbon skeleton, pyruvate or PEP depending on the decarboxylation method, is returned via gluconeogenesis and starch biosynthesis to storage starch in many species, or sucrose synthesis to storage sucrose in some species, which in turn supplies nocturnal metabolism after dusk (Dodd et al., 2002; Borland et al., 2014). The CAM cycle shares many enzymes that catalyze the key metabolic steps with the cycle of C_4_ photosynthesis (Gilman et al., 2022). For example, primary atmospheric CO_2_ assimilation is catalyzed by PPC in both CAM and C_4_ photosynthesis, and PPC activity is modulated by protein phosphorylation through PPC kinase (PPCK; Hartwell et al., 1999; Aldous et al., 2014). In the spatially separated C_4_ cycle, after decarboxylation, the three-carbon skeleton, either alanine, pyruvate, or PEP, is returned to the mesophyll cells where PPC is localized which allows the biochemical cycle to continue. In the temporally separated CAM cycle, the C_3_ acid generated due to malate decarboxylation in the light period is returned to storage starch or sucrose, which in turn must be remobilized to PEP after dusk. Nocturnal malic acid storage in the vacuole involves active import of malate via an inward rectifying voltage-gated malate channel, which could be catalyzed by a tonoplast-localized member of the *ALUMINUM-ACTIVATED MALATE TRANSPORTER* (ALMT) gene family, and export, possibly by the TONOPLAST DICARBOXYLATE TRANSPORTER (TDT), or an unknown transporter. The active transport of malate into the vacuole throughout the night is energized via the proton pumping vacuolar tonoplast adenosine triphosphatase (V-ATPase) and/ or pyrophosphatase (V-PPase; Emmerlich et al., 2003; Kovermann et al., 2007; Martinoia et al., 2007).

Storage of the carbon skeletons during the light period may occur as starch or soluble sugars, depending on the specific CAM species (Christopher and Holtum, 1998). The storage of soluble sugars to fuel nocturnal PEP supply in CAM plants has received relatively limited attention, but starch synthesis and degradation have been extensively studied in C_3_ plants, and this has led to more studies on starch synthesis and turnover in CAM species. The polymeric chains of glucose that constitute starch are synthesized in the chloroplast. ADP-glucose pyrophosphorylase (ADP-GPPase) is considered the first committed step in the starch biosynthesis pathway, and it catalyzes the conversion of glucose 1-phosphate to ADP-glucose. Starch synthases catalyze the formation of α-1,4 linkages and branching enzymes promote α-1,6-glucan linkages, while debranching enzymes selectively hydrolyze α-1,6 linkages in branched glucans (Keeling and Myers, 2010). Together with other enzymes and factors, this results in the formation of insoluble starch granules that can occupy the bulk of the volume of the chloroplast by the end of the light period.

At night, leaf starch is degraded either phosphorolytically or amylolytically, with the first steps likely being shared by both routes (Weise et al., 2006). Glucose-6-phosphate (G6P) is the final product of phosphorolytic starch degradation which is exported from the vacuole via the G6P/phosphate transporter (GPT) (Weise et al., 2006). Some CAM species have been demonstrated to degrade starch phosphorolytically rather than amylolytically (Neuhaus and Schulte, 1996; Ceusters et al., 2021).

In both CAM and C_4_, the trait evolved from the ancestral C_3_ pathway of CO_2_ assimilation during photosynthesis, in which stomata open in the light and Rubisco fixes atmospheric CO_2_ directly into the CBB cycle (Goolsby et al., 2018). Both adaptations lead to comparatively high fluxes of key metabolites and create pools of metabolites in the cell that are not present in C_3_ (Arrivault et al., 2017). In C_4_ species, conserved and repeated amino acid (AA) sequence changes have been detected in the C_4_-recruited copies of key enzymes such as PPC, and these changes are assumed to support the increased flux and metabolite pool sizes. For C_4_ plants, alignments (Paulus et al., 2013), domain swapping (Bläsing et al., 2000), and analysis of selection (Christin et al., 2007) have identified a critical alanine to serine mutation at position 780 in PPC (numbered based on the C_4_ *Flaveria* PPC sequence). Changes to NADP-ME and Rubisco AA sequences were also identified (Christin et al., 2008; Christin et al., 2009). A genome-wide scan identified three enzymes and four transporters of the C_4_ cycle as having residues under positive selection (Huang et al., 2017). The C_4_ cycle hence selects for specific changes to the AA sequences of recruited proteins, even across multiple independent origins. Some reports have suggested that particular AA changes are also associated with CAM-recruited copies of enzymes. For example, PPC was reported to carry similar changes in C_4_ and CAM species (Christin et al., 2014). Classic analysis of selection using both CAM and C_4_ species reveals AAs under selection but also questions the selection analysis as a tool due to finding a comparable amount of AAs under selection in a random set of background genes (Goolsby et al., 2018). Rubisco’s AAs have been identified as being under selection in orchids and bromeliads, some in relation to leaf δ^13^C value (Hermida-Carrera et al., 2020). The more extensive studies reported to date for specific residues occurring in the C_4_-recruited gene copies raise the question of whether or not CAM specific AA changes or changes in AA function can be identified by sampling a large and diverse set of CAM species in comparison to C_3_ species. Since CAM evolved in at least 370 genera in 38 families of plants, with a minimum of 66 independent origins of CAM, and the possibility of as many as 114 origins (Gilman et al., 2023; Holtum, 2023), it should be amenable to similar analyses compared to C_4_, which evolved independently at least 60 times (Sage et al., 2011).

Both C_4_ and CAM also require changes at regulatory levels. Transcript abundance of C_4_ cycle genes is increased several-fold (Bräutigam et al., 2011) to a degree which can be diagnostic of a “C_4_ cycle gene” vs. a non-C_4_ isoform (Christin et al., 2014; Huang et al., 2017). In the drought-inducible CAM species *Talinum triangulare* (now *T. fruticosum*), some, but not all, CAM cycle enzymes show at least one isoform with several-fold increased transcript abundance (Brilhaus et al., 2016). Similar observations were made for another inducible CAM species, *Mesembryanthemum crystallinum* (Cushman et al., 2008b). For monocot CAM species in the genus *Yucca*, some CAM cycle enzymes showed higher transcript levels in the CAM performing species as compared to a closely related C_3_ species (Heyduk et al., 2019; Heyduk et al., 2022). The C_4_ cycle spatially separates PPC and Rubisco activity and consequently requires spatial separation of activity for different enzymes and transporters (John et al., 2014). In comparison, the CAM cycle requires temporal but not spatial separation. The temporal coordination of transcript abundance for CAM is achieved by circadian clock control of many CAM-recruited gene copies (Hartwell et al., 1999; Boxall et al., 2005; Cushman et al., 2008b). In the model C_3_ species *Arabidopsis thaliana*, more than 40% of the transcriptome was found to display a diel cycle of the steady-state transcript abundance (Mockler et al., 2007; Ferrari et al., 2019). Experiments in the CAM plants *Agave americana*, *Ananas comosus*, *M. crystallinum*, *Kalanchoë fedtschenkoi* and *Sedum album* have shown that transcripts cycle but rarely show zero abundance in times where the enzyme activity should be off (Cushman et al., 2008b; Abraham et al., 2016; Sharma et al., 2017; Yang et al., 2017; Wai et al., 2019). Proteins have also been shown to cycle in their abundance in CAM *Agave americana* and *K. fedtschenkoi* (Abraham et al., 2016; Abraham et al., 2020). RNA-seq data-sets that provide a time-course of sampling at regular intervals over a 24-h light/dark cycle have allowed several studies to test whether CAM gene transcript abundances cycle robustly (JGI data release Project ID: 1155736; Abraham et al., 2016; Sharma et al., 2017; Yang et al., 2017; Wai et al., 2019). Here, we hypothesized that the obligate CAM plant *K. laxiflora* would behave in a similar manner to other, previously-studied CAM plants, and would therefore also not show binary on/off patterns for transcript and protein levels.

Both the CAM and C_4_ cycles are also associated with changes in post-translational modifications of the enzymes and transporters that catalyze key metabolic steps. For example, PPC is phosphorylated at a conserved N-terminal serine residue by its specific protein kinase, PPCK, and this changes the allosteric properties of the enzyme, including decreasing its sensitivity to feedback inhibition by malate in both CAM and C_4_ (Hartwell et al., 1999). In the CAM species *K. fedtschenkoi*, the kinase *PPCK* is transcribed and translated *de novo* each night under the control of the circadian clock, and when PPCK phosphorylates PPC in the dark, PPC becomes up to 10-fold less sensitive to inhibition by malate (Nimmo et al., 1984; Hartwell et al., 1996; Hartwell et al., 1999; Boxall et al., 2017). In C_4_ plants, *PPCK* transcription and translation is induced by light leading to PPCK activity that phosphorylates PPC and also decreases its sensitivity to inhibition by malate (Hartwell et al., 1999; Aldous et al., 2014).

In this manuscript, we analyzed the obligate CAM species *K. laxiflora* in detail, embedded in the context of available CAM genome sequences. First, we tested in *K. laxiflora* whether the diel cycles of transcript and protein abundance in CAM-performing mature leaves were sufficient for the expected on/off patterns of regulation of carboxylation/decarboxylation pathway steps. Secondly, we analyzed the available AA sequences for core CAM pathway enzymes and metabolite transporters and their regulatory partners to determine whether the trait is associated with specific changes to the AA sequence of these proteins. Lastly, we tested whether post-translational modifications (PTMs) are involved more widely in regulating metabolic fluxes over each 24-h light/dark cycle in CAM plants, beyond the known regulation of enzymes such as PPC and PPDK. The results revealed that CAM gene transcript levels and the abundance of the encoded proteins did not show a binary on/off pattern between light and dark. We also discovered that the candidate CAM-recruited gene family members across many CAM species shared AA changes that were not shared by the orthologs and paralogs in C_3_ species. These CAM-associated conserved changes in enzyme AA sequences were over-represented in enzymes associated with starch metabolism. Finally, several phosphoproteome changes were detected in the CAM leaves of *K. laxiflora,* which revealed that PTMs on CAM proteins that have not been reported previously may be relevant to the regulation of the 24-h light/dark cycle of CAM.

## Methods

### Plant material

*Kalanchoë laxiflora* plants (Accession 2014091.4; source information in supplemental table 1) were grown in a greenhouse in long day conditions (16-h-light/8-h-dark; 20°C in the light and 18°C in the dark) for about 22-weeks in 10 cm pots (compost mix consisting of half volume sand “Wesersand” and half volume “Fruhsdorfer Einheitserde T”). Leaf pairs (LP) 6-7, where LP1 are the youngest, newly emerging leaves either side of the shoot apical meristem, were harvested and snap-frozen in liquid nitrogen in the middle of the light and dark period (corresponding to zeitgeber time (zt) zt8 and zt20, respectively) in five replicates for proteomic analysis and in three replicates for RNA-seq analysis. Plant tissue was ground under liquid nitrogen to a fine powder and stored at - 80°C until use. Five further CAM species (Source information in Supplemental Table 1) were selected from the Bielefeld University greenhouse collection, namely *Euphorbia abyssinica*, *Pachypodium lamerei*, *Xerosicyos danguyi*, *Dendrobium delicatum* and *Ariocarpus trigonus* var. *elongatus*. For these species, plant material was sampled at plus/minus two hours of the plants mid-day to avoid diurnal changes between samples and snap-frozen in liquid nitrogen.

### RNA-seq

After grinding in liquid nitrogen, RNA was extracted using Qiagen RNeasy Plant Mini Kit according to manufacturer’s instructions with addition of 13.5 µL PEG20,000 (50 mg mL^-1^ stock) to 450 µL RLC buffer. Sequencing libraries were constructed using the Illumina TruSeq library kit according to the manufacturer’s instructions and sequenced to a depth of 5.3 to 6.7 million reads on the Illumina NextSeq500 for *E. abyssinica*, *P. lamerei*, *A. trigonus* var. *elongatus*, *X. danguyi*, and *D. delicatum*, and to a depth of 7.6 to 8.8 million reads on the Illumina NextSeq1000 for *K. laxiflora*. For quantitative analysis, the *K. laxiflora* reads were mapped against the reference with kallisto 0.44.0 (Bray et al., 2016) with the parameters set to -b 100, -l 180, -s 20 and using the --single flag. Differentially abundant transcripts were detected with edgeR using the classic mode (Robinson et al., 2009). Transcripts were considered significantly differentially abundant if FDR-adjusted p-value (q-value) < 0.01. Gene Ontology (GO) enrichments in groups were determined with TopGO using a classic Fisher’s exact test without adjustment (Alexa and Rahnenfuhrer, 2024) after GO annotation transfer via protein blast from *A. thaliana*. For the qualitative analysis, the sequencing reads for *E. abyssinica*, *P. lamerei*, *X. danguyi*, *D. delicatum* and *A. trigonus elongatus* were trimmed by Trimmomatic (settings: LEADING:3 TRAILING:3 SLIDINGWINDOW:4:15 MINLEN:3; Bolger et al., 2014) and used for *de novo* assembly with Trinity v2.8.4 (Haas et al., 2013) using default settings (--seqType fq --full_cleanup --normalize_by_read_set) and transcript sequences were translated to amino acid sequences using TransDecoder.LongOrfs (https://github.com/TransDecoder/TransDecoder.git).

RNAseq data generated during this study is available at NCBI SRA under BioProject ID PRJNA1172678.

### Proteins – AA Conservation – Data

Protein FASTA files for *Kalanchoë fedtschenkoi*, *Eucalyptus grandis*, *Citrus clementina*, *Brassica oleracea*, *Theobroma cacao*, *Ricinus communis*, *Gylcine max*, *Arachis hypogaea*, *Trifolium pratense*, *Medicago truncatula*, *Fragaria vesca*, *Beta vulgaris*, *Helianthus annuus*, *Mimulus guttatus*, *Solanum lycopersicum*, *Brachypodium distachyon*, *Hordeum vulgare*, *Oryza sativa*, *Dioscorea alata*, *Amborella trichopoda*, and *Spirodela polyrhiza* were downloaded from JGI Phytozome v13 (Goodstein et al., 2012). Protein FASTA files for *Vanilla planifolia*, *Hylocereus undatus* (also known as *Selenicereus undatus*; Liu et al., 2021), *Simmondsia chinensis*, *Cucumis sativus* L., *Asparagus officinalis*, *Allium sativum*, and *Actinidia chinensis* were downloaded from PLAZA (Van Bel et al., 2022). The *Sedum album* protein FASTA file was obtained from CoGe (https://genomevolution.org/coge/OrganismView.pl?oid=38427 version V3). The protein FASTA file for *Arabidopsis thaliana* was downloaded from TAIR (TAIR10_pep_20110103_representative_gene_model_updated). Protein FASTA files for *Apostasia shenzhenica*, *Dendrobium catenatum*, and *Phalaenopsis equestris* were obtained from the OrchidBase (Fu et al., 2011). Publicly available RNA-seq data was downloaded and used for *de novo* assembly with Trinity v2.8.4 as described above (Haas et al., 2013) for *Cymbidium atropurpureum*, *Dendrobium terminale*, *Cymbidium mannii*, *Hylocereus polyrhizus*, *Lophophora williamsii*, *Opuntia ficus-indica*, *Talinum fruticosum*, and *Yucca aloifolia*. For the full list of references for genome data and SRA (sequence read archive) numbers see supplemental table 3. We used TransDecoder (https://github.com/TransDecoder/) to obtain protein sequences where only transcriptome data was available, as described above.

### AA Conservation and Changes

OrthoFinder v2.3.7 (Emms and Kelly, 2015) connected the orthologues of the 47 species in the analysis. For each orthogroup (OG) containing a CAM protein as identified by the Arabidopsis ID (for a list see Schiller and Bräutigam, 2021), ClustalO v1.2.4 (Sievers et al., 2011) aligned all protein sequences within the respective OG using the default settings with only the longest isoform in the OG retained for the data assembled with Trinity. The OG containing AMY3 was split manually using the phylogenetic tree generated by ClustalO, because it contained not only Alpha Amylase 3 (AMY3), but also AMY1 and AMY2.

A custom python script was used to determine amino acid positions that were conserved across the C_3_ species, but displayed changes in CAM species. Conservation of a position was defined as 80% identity across all C_3_ sequences, while functional conservation of a position was defined as 80% identity in AA groups (non-polar, polar, acidic, and basic) across all C_3_ sequences. Since CAM is not a binary trait and since different CAM species may use alternative paths to the same end, we did not expect binary changes in the conservation pattern. For every conserved position we therefore determined whether the CAM sequences were different from the conserved AA (or group) in at least half of the sequences.

### Proteomics Analyses

Proteins were extracted from powdered leaves with lysis buffer (4% (w/v) SDS, 100 mM Tris/HCl pH 7.6, 2.5 : 1, vol/sample weight) and prepared for MS analysis following the SP3 protocol (Hughes et al., 2019), but with the modification that 50 mM TCEP and 55 mM CAA were used for reduction and alkylation. Aliquots of 50 µg peptides were fractionated for full proteome analysis (Deng et al., 2021). For phosphopeptide enrichment, an input of 500 µg was used following a previously published procedure (Nakagami, 2014). LC-MS/MS analysis was performed by using an EASY-nLC 1200 (Thermo Fisher) coupled to an Exploris 480 mass spectrometer (Thermo Fisher). Separation of peptides was performed on 20 cm frit-less silica emitters (CoAnn Technologies, 0.75 µm inner diameter), packed in-house with reversed-phase ReproSil-Pur C_18_ AQ 1.9 µm resin (Dr. Maisch). The column was constantly kept at 50 °C. Peptides were eluted in 75 min applying a segmented linear gradient of 0 % to 98 % solvent B (solvent A 0 % ACN, 0.1 % FA; solvent B 80 % ACN, 0.1 % FA) at a flow-rate of 300 nL/min. Mass spectra were acquired in data-dependent acquisition mode. MS^1^ scans were acquired at an Orbitrap Resolution of 120,000 with a Scan Range (m/z) of 280-1500, a maximum injection time of 100 ms, and a normalized AGC Target of 300 %. For fragmentation, only precursors with charge states 2-6 were considered. Up to 20 Dependent Scans were taken. For dynamic exclusion, the duration was set to 40 sec and a mass tolerance of +/- 10 ppm. The isolation window was set to 1.6 m/z with no offset. A normalized collision energy of 30 was used. MS^2^ scans were taken at an Orbitrap Resolution of 15,000, with a fixed First Mass (m/z) = 100. Maximum injection time was 150 ms and a normalized AGC Target 5%. MS raw data were analyzed using MaxQuant (MQ) 2.1.3.0 and searched against a custom in house *K. laxiflora* protein FASTA and a list of common contaminants and a reverse decoy database was included. All raw files from the proteome and phosphoproteome were searched within one run using different parameter groups and separated fractions. The ‘match between runs’ feature was enabled. Carbamidomethylation was used as fixed modification and oxidation of methionine and N-terminal protein acetylation were set as variable modifications. In addition, phosphorylation of serine, threonine and tyrosine were searched as variable modifications in the respective parameter group for the phosphopeptide-enriched samples. Label-free quantification (LFQ) and intensity based absolute quantification (iBAQ) were enabled for the parameter group of full proteome runs. Raw and MQ txt files, as well as the custom protein FASTA file are available via JPost identifier JPST003223. Up until publication of the manuscript, the following credentials can be used to access the data URL https://repository.jpostdb.org/preview/1787248601669f60e52d36d Access key 2604

Data was further analyzed with LIMMA (Ritchie et al., 2015) after filtering for quantification in at least 3 out of 5 replicates in one condition. Protein groups that were only detected in one condition with missing data in all five replicates of the other condition are reported separately. Protein groups that were detected in both timepoints were treated as follows: All missing values were omitted from statistical analysis. LIMMA was applied on the LFQ values and log_2_ fold changes were calculated by log_2_((mean_day + 1)/(mean_night + 1)). P-values were adjusted with Benjamini-Hochberg adjustment (Benjamini and Hochberg, 1995). Protein groups were considered significantly differentially abundant if adjusted p-value < 0.05. Gene Ontology enrichments were determined for protein groups that were significantly more abundant at night together with protein groups that were only detected at night and for protein groups that were significantly more abundant during the day together with those that were only detected during the day. Enrichments were determined with TopGO using classic Fisher’s exact test (Alexa and Rahnenfuhrer, 2024) after GO annotation via protein blast from *A. thaliana*.

Phosphopeptides that were quantified at both timepoints with at least 3 out of 5 replicates of one condition identified were analyzed for differential abundance using LIMMA (Ritchie et al., 2015). LIMMA was applied on the intensity-values and the log_2_ fold changes were calculated as follows: log_2_((mean_day + 1)/(mean_night + 1)). P-values were adjusted by Benjamini-Hochberg adjustment (Benjamini and Hochberg, 1995) and phosphopeptides were considered significantly differentially abundant if adjusted p-value < 0.05. Phosphorylated peptides that were only detected in one condition are reported separately.

### Conservation of potential PTM sites

For analysis of the conservation of amino acids that are putative targets to PTM, we used the same OGs and alignments of 47 angiosperm species as described before. No distinction between C_3_ and CAM photosynthesis type was made because the aim was to identify the regulatory architecture, which was predicted to be conserved between CAM and C_3_ plants.

The 78 *A. thaliana* orthologues of CAM proteins (Schiller and Bräutigam, 2021; supplemental table 4) were analyzed for the occurrence and positions of PTMs (phosphorylation, acetylation, S-nitrosylation) in *A. thaliana* using the database FAT-PTM (Cruz et al., 2019). To test their potential for regulation in CAM species, the conservation of the amino acid (AA) at the respective position in the alignment across the orthologues was determined. The AA was considered conserved if it was identical to the *A. thaliana* AA in at least 70% of the available sequences.

## Results

### Transcript and protein abundances in general

Three replicate mature leaf samples were collected in the middle of the 16-h-light period and middle of the 8-h-dark period (zt8 and zt20, respectively) from well-established plants of the obligate CAM species *K. laxiflora,* and transcript abundance was analyzed via RNA-seq (Figure 1a). Of all transcripts, 71.8% (19,941 of 27,768) were detected (transcripts per million (tpm) >1 in at least 2/3 replicates in at least one condition). At zt8, 69.1% (19,192/27,768) of transcripts were detected, and at zt20, 70.1% (19,465/27,768) of all transcripts were detected.

**Figure 1:**
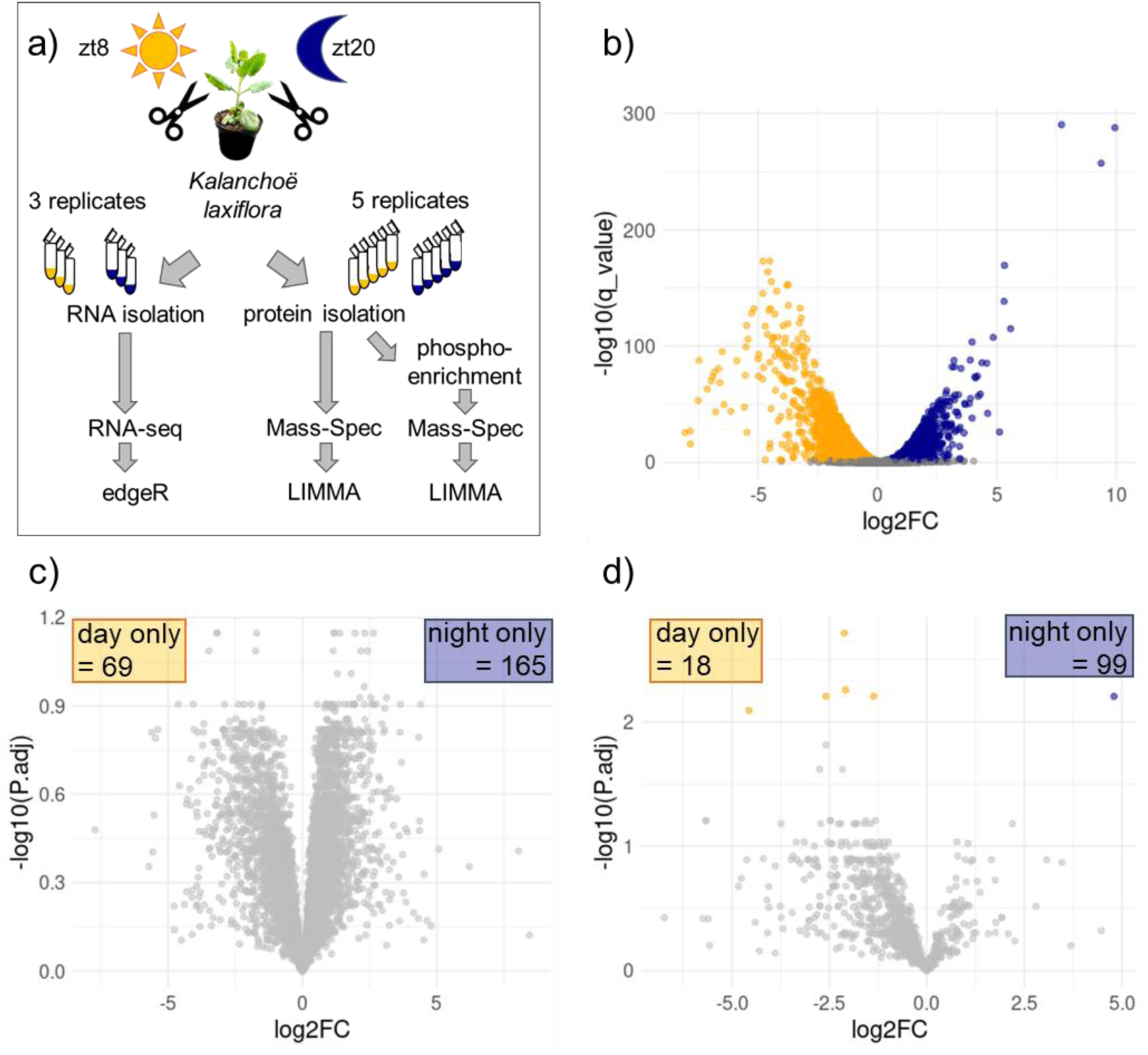
Transcriptome and proteome analyses of Kalanchoë laxiflora leaf material from day and night conditions. a) Workflow for analysis of differential transcript and protein expression in K. laxiflora grown in the greenhouse under 16h light 8h dark conditions. zt = zeitgeber time. b) Volcano plot of the differentially expressed transcripts between day and night in K. laxiflora LP6. Number of quantified items: 27,768. Orange: significantly (q-value<0.05) downregulated at night Blue: significantly (q-value<0.05) upregulated at night Grey: not significantly differentially expressed. c) Volcano Plot showing differentially abundant protein groups between mid-day and mid-night in K. laxiflora (quantified by LC-MS/MS analyses). Number of quantified items: 5,241. Grey: not significantly differentially expressed. Numbers for day only and night only indicate how many protein groups were only detected at one timepoint with missing values for all replicates of the other timepoint d) Volcano plot showing differentially abundant phosphorylated peptides between mid-day and mid-night in K. laxiflora (quantified by LC-MS/MS analyses). Number of quantified items: 794. Orange: significantly (q-value<0.05) downregulated at night Blue: significantly (q-value<0.05) upregulated at night Grey: not significantly differentially expressed. Numbers for day only and night only indicate how many phospho-peptides were only detected at one timepoint with missing values for all replicates of the other timepoint.

Differential expression analysis resulted in 6,247 significantly (q-value < 0.01) altered transcripts between the two time points. Of these significantly differentially expressed genes, 3,539 have a log_2_FC between -1 and 1 (Figure 1b). There were 3,167 significantly downregulated genes at night and 3,080 significantly upregulated genes at night. Transcripts downregulated at night enrich in 39 GO Terms. The most significant enrichment (p = 1.4e-5) is found for the term photosynthesis. Other enriched GO terms are related to biosynthetic and metabolic processes as well as response to water and light stimulus (supplemental table 6). During the night, upregulated transcripts enrich in GO terms related to responses to various stimuli like ethylene, abiotic stimulus, osmotic stress, response to fungus (supplemental table 5).

Five replicate mature leaf samples were also harvested at zt8 and zt20 and used for protein abundance analysis via label free quantitative mass spectrometry (Figure 1a). After filtering (identification in at least 3/5 replicates in at least one condition), 5,475 of 27,768 potentially possible protein groups were called as identified. At mid-day, 4,915 protein groups were identified in at least 3/5 replicates and at mid-night, 5,191 protein groups were identified in at least 3/5 replicates (Figure 1c). When considering only proteins that were detected at both timepoints, none of them were significantly differentially abundant between mid-day and mid-night (q-value < 0.05) indicating a fairly stable proteome in *K. laxiflora* at these two time points that was independent of the time of sampling in the light and dark (Figure 1c). One hundred and sixty-five protein groups were detected exclusively at zt20, whereas 69 protein groups were only detected at zt8 (Figure 1c). GO enrichment analyses of the significantly upregulated proteins at night, including those exclusively detected at night, resulted in 18 significantly enriched GO terms (p-value < 0.05), many of which are related to nucleic acid metabolic processes (supplemental table 8). Combining the upregulated protein groups during the day and those exclusively detected at zt8 resulted in two significantly enriched GO terms, ion transport and system development (supplemental table 7).

### Transcript and protein abundances of CAM genes

To highlight the potential CAM-associated transcripts and encoded proteins that changed in abundance between the light and the dark, 84 members of gene families that are predicted to include the core CAM pathway genes were defined for *K. laxiflora* based on the list of CAM proteins in Schiller and Bräutigam (2021). Genes were identified due to orthology and grouped with the CAM pathway (Figure 2, note the log-scale). In *K. laxiflora*, CAM transcripts were detected at abundances between 0.1 and 5,626 transcripts per million with GLYCERALDEHYDE-3-PHOSPHATE DEHYDROGENASE C2 (*Kl*GAPC2) being the most abundant and *Kl*PPC4 being lowest (Figure 2a). Of the 84 potentially CAM-associated genes, 42 had transcript levels that were significantly differentially abundant between zt8 and zt20, of which 20 had a |log_2_FC| > 1 (Figure 2a). Assuming that gene transcript abundance in mature, full-CAM leaves is a reliable predictor of CAM pathway membership for the encoded protein, as it is accepted for C4, specific CAM-associated gene family members were assigned for each enzymatic and metabolite transport step of the pathway, along with the relevant known regulatory proteins such as PPCK.

**Figure 2:**
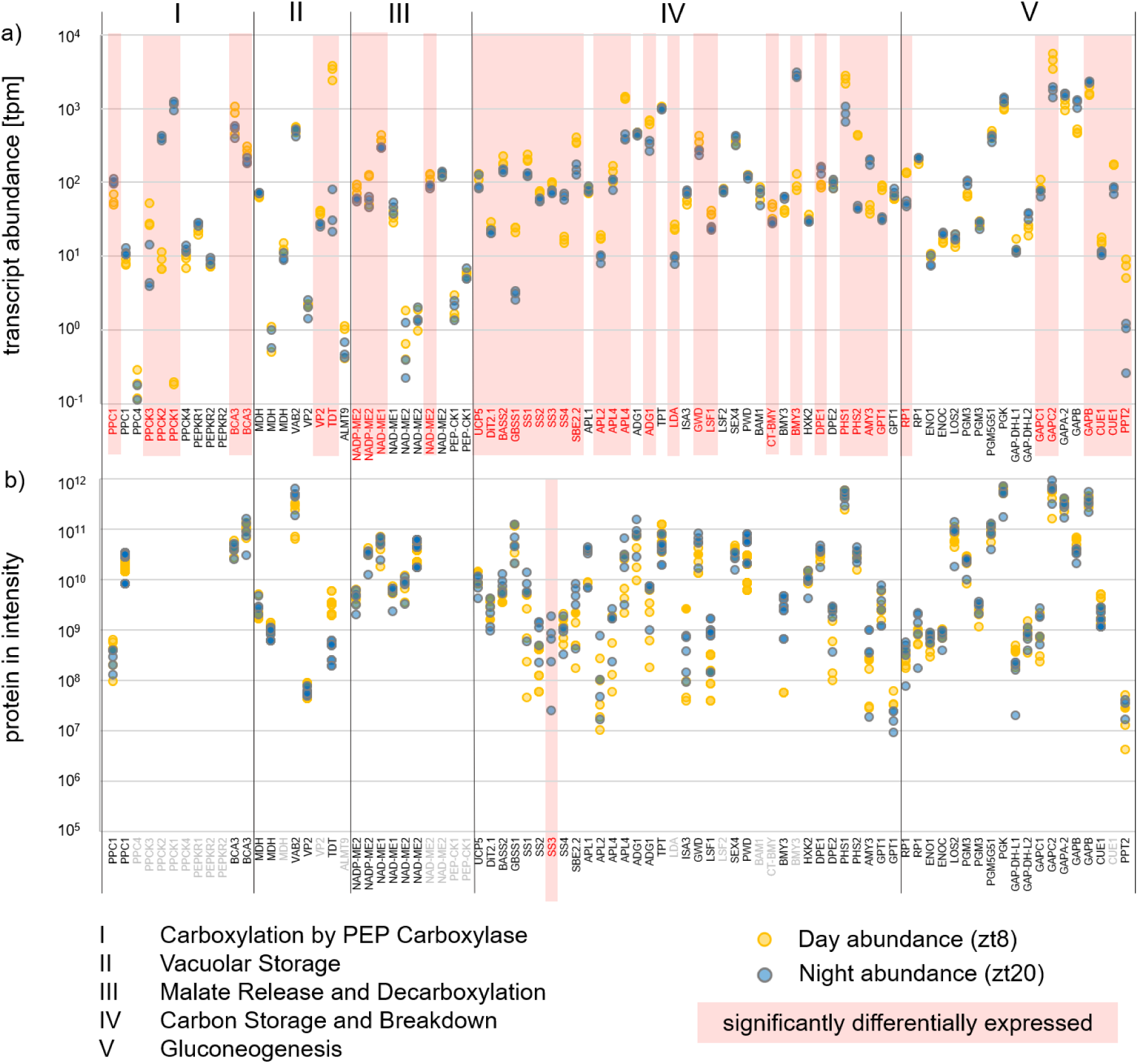
Transcriptome and proteome analysis of gene products involved in CAM-metabolism. a) tpm values for CAM transcripts in K. laxiflora at mid-day = zt8 (yellow) and mid-night = zt20 (blue), naming based on A. thaliana orthologues. Scale is logarithmic. Red names indicate significantly differentially expressed transcripts (q-value < 0.01). b) LFQs for CAM proteins in K. laxiflora at mid-night (blue) and mid-day (yellow). Note that y-axis is logarithmic and starts at 10^5^. Names based on A. thaliana orthologues. Red names indicate differentially abundant proteins (q-value <0.05 or only detected at one timepoint), grey names indicate proteins that were not identified at both timepoints. Gene IDs are sorted based on their protein’s involvement in the different groups I: Carboxylation by PEP carboxylase; II: Vacuolar Storage; III: Malate Release and Decarboxylation; IV: Carbon Storage and Breakdown; V: Gluconeogenesis. Abbreviations: ADG: ADP glucose pyrophosphorylase; ALMT: Aluminum-activated malate transporter; AMY: Alpha amylase; APL: Glucose-1-Phosphate adenylyltransferase; BASS: Bile-acid sodium symporter; CA: carbonic anhydrase; DiT: Dicarboxylate translocator; DPE: Disproportionating enzyme; ENO: Enolase; GBSS: Granule-bound starch synthase; GPT: G6P Phosphate transporter; HXK: hexokinase; MDH: malate dehydrogenase; NAD(P)-ME: NAD(P) dependent malic enzyme; PEP-CK: Phosphoenolpyruvate Carboxykinase; PGM: Phosphoglucomutase; PHS: Starch phosphorylase; PPase: Pyrophosphatase; PPC: PEP carboxylase; PPCK: PPC kinase; PPDK: Pyruvate, orthophosphate dikinase; PPT: PEP/Phosphate translocator; RP: regulatory protein; SBE: Starch branching enzyme; SS: Starch synthase; TDT: Tonoplast dicarboxylate transporter; TPT: Triosephosphate/phosphate translocator

For some CAM genes *sensu stricto*, a CAM-associated member of the gene family could be assigned based on the RNA-seq data presented here. Three *K. laxiflora* genes were assigned as *PPC* orthologues. The *Kl*PPC4 orthologue had an abundance of less than one transcript per million in all replicates at both timepoints and one *PPC* orthologue had tpm values of ∼10 across all replicates at both timepoints. The last *PPC1* orthologue (KlGene020155-RA) can be assigned as the CAM-isoform according to transcript abundance. With tpms ranging from ∼50 during the day to about ∼100 during the night it is the highest expressed PPC transcript in *K. laxiflora*. The difference in day/night transcript abundance is significant with a log_2_FC of 0.8 (adjusted p value = 0.00045). Both analyzed *β-CA* orthologues were downregulated at night with tpm ranges between 180-310 and 400-1000 for the orthologues, respectively. In the group of CAM genes encoding for proteins involved in metabolite transport of malate into and out of the vacuole, *TDT* was strongly down-regulated at night with the second highest log_2_FC of -6. The mean tpm was 44 at zt20 and 3,226 at zt8 for *TDT*.

Decarboxylation in *K. laxiflora* is mediated by ME (Dever et al., 2015). One orthologue of *NAD-ME1* alpha subunit was detected at a higher transcript abundance compared to all others with a mean tpm of 395 at zt8 and 295 at zt20, resulting in a log_2_FC of -0.4 (KlGene001496-RA; Figure 2a). Three additional *Kl*ME orthologues (two NADP-ME and one NAD-ME beta orthologues) were significantly differentially abundant with log_2_FC between - 0.4 and -0.8 and tpms between 28 and 141. Other significantly differentially abundant transcripts in the group of malate release and decarboxylation include the dicarboxylate carrier *UCP5*, dicarboxylate translocator *DIT2*, and the Bile-Acid Sodium Symporter *BASS2*, all of which are slightly upregulated during the day (log_2_FC between 0.3 and 0.4) with tpm values in the range of 20-29 up to 138-230.

Twenty transcript isoforms of proteins involved in starch synthesis and breakdown had significantly differential abundance (Figure 2a). These were Granule Bound Starch Synthase 1 (GBSS1), Starch Synthase 1/2/3 (SS1/2/3), Starch Branching Enzyme 2 (SBE2), Glucose-1-Phosphate Adenylyltransferase 2/4 (APL2/4), ADP Glucose Pyrophosphorylase (ADG1), Limit Dextrinase (LDA), Glucan Water Dikinase (GWD), Like Sex Four 1 (LSF1), Cytosolic Beta-Amylase (CT-BMY), Starch Phosphorylase 1/2 (PHS1/2), and GPT1 being significantly upregulated during the day (with log_2_FC between -0.35 and -3.2) and SS4, Beta Amylase 3 (BMY3), Disproportionating Enzyme 1 (DPE1), and Alpha Amylase (AMY) being significantly upregulated at night (with log_2_FC between 0.73 and 4.8; compare supplemental table 4 and Figure 2a). In the group of proteins involved in gluconeogenesis, transcripts of GAPC1/2, PEP/phosphate translocators CUE1 and PPT2 were significantly downregulated at night and one isoform of GAPB was significantly upregulated at night (Figure 2a, supplemental Table 4). For PPDK regulatory protein (RP), one isoform (KlGene013132-RA) is significantly differentially abundant (log_2_FC -1.4) with tpms between 50 (night) and 135 (day). However, absolute transcript abundance was lower than for the second RP orthologue that was detected at ∼ 200 tpm at both timepoints (Figure 2a).

The only gene with true binary presence/absence variation among the CAM genes was *PPCK1* (KlGene014151-RA). This ortholog of the previously characterized *K. fedtschenkoi* CAM-associated *PPCK1* was detected at tpms below one during the day and above 1000 at night, and had a log_2_FC of 10 and could thus be assigned as the CAM-associated *PPCK1* in *K. laxiflora* based on transcript abundance (Figure 2a). The other *PPCK* gene family members were either not differentially abundant between the two time points measured, had a lower tpm span (from ∼8 at zt8 to ∼400 at zt20), or were differentially abundant in the other direction (∼35 tpm at zt8 and ∼7 tpm at zt20).

Among the 84 genes belonging to the gene families of the core set of CAM-associated genes in *K. laxiflora*, 62 were identified within the protein abundance data (after filtering); none of them were significantly differentially abundant between zt8 and zt20 (adjusted p-value <0.05). SS3 was only identified at zt20. The other CAM proteins were either not detected at both timepoints or had a similar abundance in both the dark and light samples (Figure 2b). *PPCK1* had highly differential transcript abundance, being low at zt8 and over 1000-fold higher at zt20, but the protein was not detectable at both timepoints. No peptides were identified for both PEP-CK isoforms, ALMT9, two NAD-ME2 beta isoforms and one isoform of CUE1, Vacuolar H^+^ Pyrophosphatase 2 (VP2) and MDH. All of these proteins also had low transcript abundance at both time points (Figure 2a). Although LDA, Like SEX Four 2 (LSF2), Beta Amylase 1 (BAM1), Cytosolic Beta Amylase (CT-BMY), and one isoform of Beta Amylase 3 (BMY3) were detected at the transcript level (Figure 2a), no peptides were detected for these proteins.

In summary, only the gene encoding the PPC-specific protein kinase, PPCK, displayed a binary on/off of its transcript level between the dark (on) and light (off) time points. The genes encoding proteins that catalyze the flux through the CAM pathway were found to have differential transcript but not protein abundance, and did not show a binary on/off pattern at the mid-day and mid-night time points sampled here.

### Conserved amino acid sequence changes within CAM proteins across CAM species

To examine potential adaptations associated with CAM at the protein sequence level, we collected publicly available protein sequence data for 15 CAM species and 27 C_3_ species. To cover taxonomic orders within the Euasterids and Fabids with known CAM origins, and to extend the available data in the CAM-rich orders Asparagales and Caryophyllales, we sequenced RNA of five CAM species and assembled their transcriptomes *de novo*. RNA from leaves of *Pachypodium lamerei* (Figure 3; Apocyanaceae, Gentianales) was sequenced. Within the Fabids, three orders with known CAM origins have been reported, of which we covered two with *de novo* transcriptome sequencing and assembly for *Euphorbia abyssinica* (Figure 3b; Euphorbiaceae, Malpighiales) and *Xerosicyos danguyi* (Figure 3c; Cucurbitaceae, Cucurbitales). Within the Caryophyllales, we added *Ariocarpus trigonus* var. *elongatus* (Figure 3d; Cactaceae, Caryophyllales) to extend our sampling of this CAM-rich order, and for monocots we extended our sampling with a *de novo* transcriptome assembly for leaves of *Dendrobium delicatum* (Figure 3e; Orchidaceae, Asparagales).

**Figure 3:**
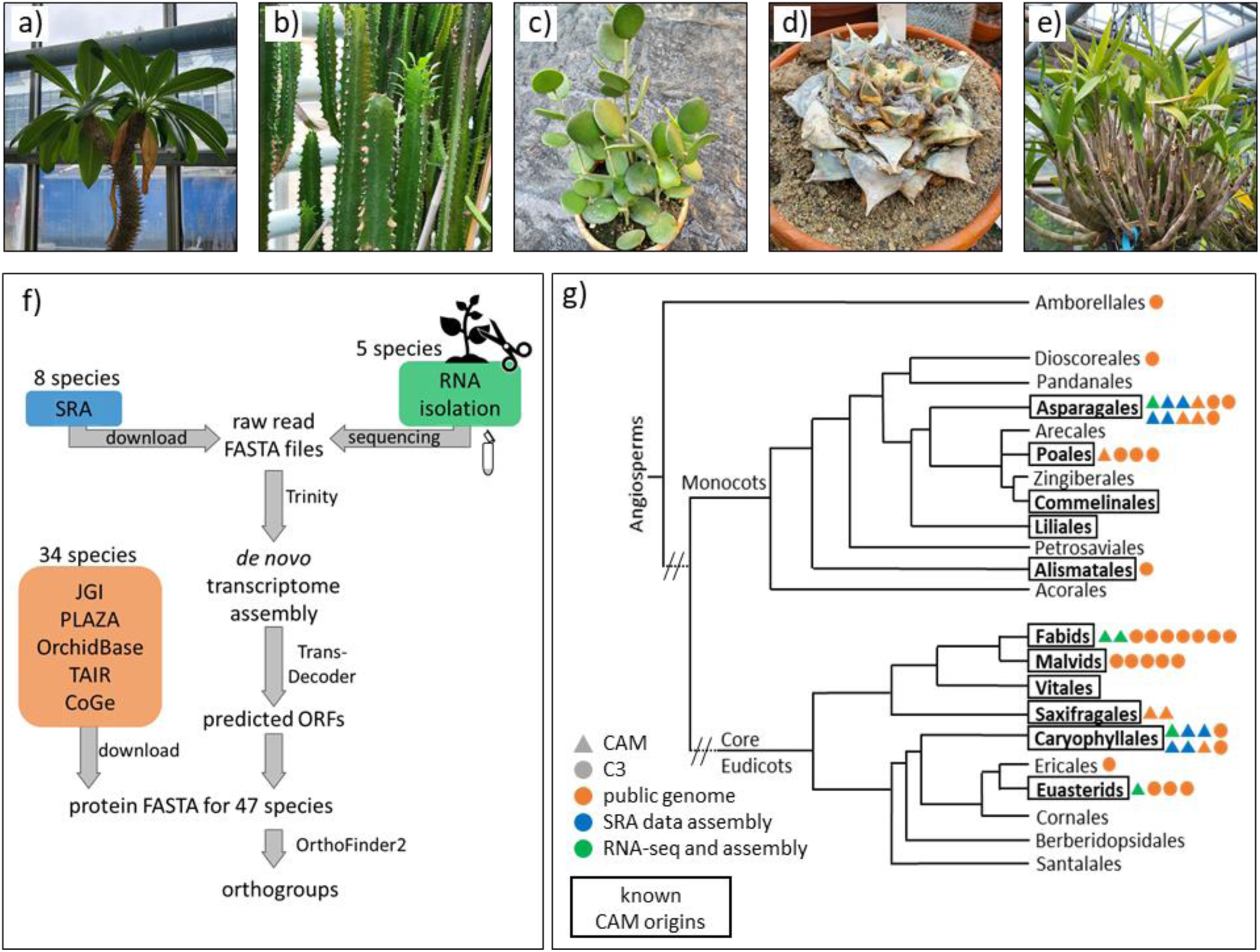
Overview of workflow and species for comparative analyses. a) Pachypodium lamerei b) Euphorbia abyssinica c) Xerosicyos danguyi d) Ariocarpus trigonus var. elongatus e) Dendrobium delicatum f) workflow for data generation and collection g) tree with used plant species marked by colored shapes. Color is based on where the data comes from: green – sampled from the greenhouse, sequenced and de novo assembly; blue – transcriptome data from the SRA for de novo transcriptome assembly; orange – protein FASTA files downloaded from databases such as JGI, PLAZA, OrchidBase, TAIR, and CoGe. Shape is based on photosynthetic type of sampled species: circles – C3; triangles – CAM. Boxes indicate known CAM origins.

Overall, the 20 CAM species sampled for their proteomes are distributed across seven orders. Within monocots, we sampled nine species in three families of two orders, and for dicots we sampled eleven species from six families in five orders. The 27 C_3_ species selected for comparative analysis against the CAM species are distributed across 18 orders. Within monocots, eight species represented six families in four orders, and for dicots, 18 species provided coverage of 14 families in 13 orders, and one species provided representation of the basal angiosperm clade Amborellales. The CAM species sampled cover two of five orders with known CAM origins within the monocots and five of ten orders with CAM origins within the core eudicots (Figure 3g). For five of the seven orders sampled with known CAM origins, we analyzed at least one CAM and one C_3_ species.

Orthologues for the 78 candidate CAM proteins in *A. thaliana* (Schiller and Bräutigam, 2021) were grouped into 56 OGs by OrthoFinder2 and orthologues were aligned (Figure 4a). Some of the CAM proteins were grouped in the same OG resulting in fewer OGs than candidate CAM proteins. For example, *At*APL1, *At*APL2, and *At*APL4 were grouped together. Two different types of changes were tested for, (i) a radical change in which functional groups change and (ii) a conservative change in which AAs change within the same group (Figure 4a). Changes in the AA sequence were allowed to be fuzzy because CAM is not a binary trait and alternative branches of the CAM pathway exist. For decarboxylation, ME or PEP-CK, and in carbon storage, starch or soluble sugars can be used. Furthermore, for many species we cannot determine the CAM recruited gene copy or copies and therefore analyzed all sequences within the orthogroups. We show in detail two examples of 154 observed changes. The disproportionating enzyme 2 (DPE2) protein involved in starch breakdown in the dark carries a conserved glutamate at position 1058 across the C_3_ species that changed to the non-polar glycine in 15 out of 23 CAM sequences resulting in a radical change (Figure 4b). Another enzyme in starch breakdown, alpha amylase 3 (AMY3), had a conserved glutamine at position 668 amongst C_3_ species, which was changed to a basic histidine or arginine in 22 out of 32 CAM sequences (Figure 4c). Across all putative CAM proteins analyzed here, a total of 154 positions were detected with fuzzily conserved changes between C_3_ and CAM species. Of these, 75 were classified as a conservative change (change within the AA group), and 79 as radical change (across groups). Out of the 56 alignments of CAM candidate proteins tested, 31 had at least one AA position with a fuzzily conserved and therefore putatively adaptive change (Figure 4d).

**Figure 4:**
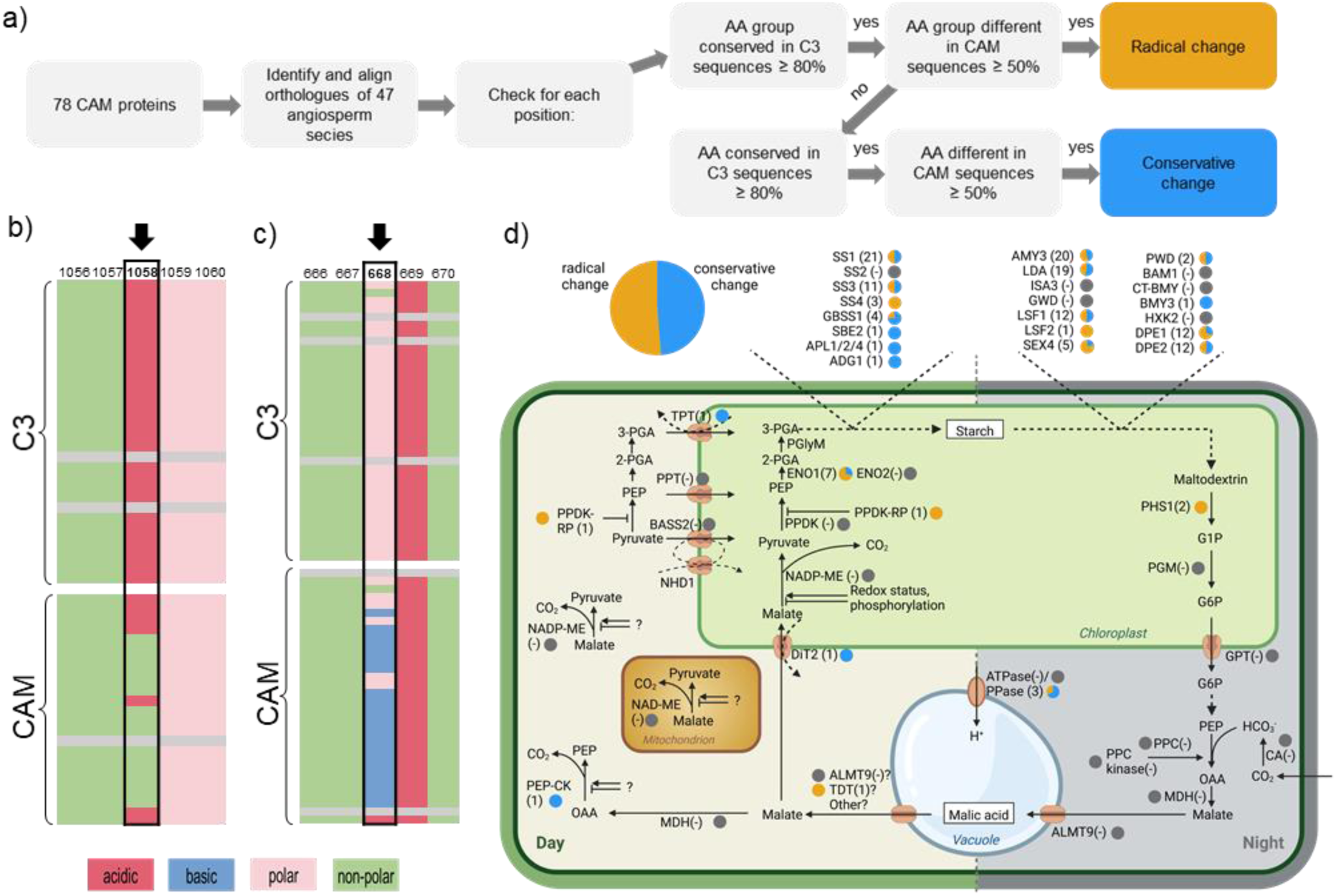
Putative adaptive changes in CAM proteins. (a) Workflow of AA position analysis. (b) and (c) examples for positions with radical changes ± 2 surrounding amino acids for DPE2 (b) and AMY3 (c) with amino acid groups color-coded (pink = polar, red = acidic, green= non-polar, blue=basic). Position with detected fuzzily conserved change is marked by a black arrow. For details see supplemental dataset 1. (d) CAM pathway with conserved putative adaptive changes. Number in brackets gives the total number of positions with changes between C3 and CAM. Little pie charts indicate the proportion of the positions that have radical (yellow) or conservative (blue) changes. Grey pie charts indicate that no fuzzily conserved change was found for the analyzed orthologues. Abbreviations: ADG: ADP glucose pyrophosphorylase; ALMT: Aluminum-activated malate transporter; AMY: Alpha amylase; APL: Glucose-1-Phosphate adenylyltransferase; BASS: Bile-acid sodium symporter; CA: carbonic anhydrase; DiT: Dicarboxylate translocator; DPE: Disproportionating enzyme; ENO: Enolase; G1P: Glucose-1-Phosphate; G6P: Glucose-6-Phosphate; GBSS: Granule-bound starch synthase; GPT: G6P Phosphate transporter; HXK: hexokinase; MDH: malate dehydrogenase; NAD(P)-ME: NAD(P) dependent malic enzyme; NHD: Sodium hydrogen antiporter; OAA: oxaloacetate; PEP: Phosphoenolpyruvate; PEP-CK: PEP carboxykinase; PGA: Phosphoglycerate; PGlyM: Phosphoglycerate mutase; PGM: Phosphoglucomutase; PHS: Starch phosphorylase; PPase: Pyrophosphatase; PPC: PEP carboxylase; PPCK: PPC kinase; PPDK: Pyruvate, orthophosphate dikinase; PPT: PEP/Phosphate translocator; RP: regulatory protein; SBE: Starch branching enzyme; SS: Starch synthase; TDT: Tonoplast dicarboxylate transporter; TPT: Triosephosphate/phosphate translocator

Examination of all candidate CAM proteins showed that 128 out of 154 fuzzily conserved detected changes among CAM pathway proteins were found in the starch synthesis and degradation pathway (Figure 4d). Twenty-one of the thirty analyzed proteins involved in these pathways had at least one position with a change between C_3_ and CAM. SS1, SS3, AMY3, LDA, DPE1 and DPE2 were found to have each accumulated more than ten fuzzily conserved, potentially adaptive changes in their protein sequences (Figure 4d). The plastid-localized PHS1 yielded two radical changes, which both occurred in the same subset of CAM species, namely the CAM species belonging to the Orchidaceae as well as Saxifragales (supplemental dataset 24). The cytosol-localized PHS2 did not yield any fuzzily conserved changes between C3 and CAM species. The cytosolic enolase (ENOC) had one conserved AA change (supplemental dataset 12). In comparison to that, the chloroplast-localized enolase had seven positions with fuzzily conserved changes (ENO1; Figure 4d, supplemental dataset 13). The core CAM enzymes *sensu stricto* (CA, PPC, PPCK, MDH, ME, PPDK) did not yield any fuzzily conserved changes with the exception of one candidate position in PEP-CK with a conservative change from a serine in C_3_ species to a cysteine in half of the CAM species (Figure 4d; supplemental dataset 21).

Changes in sequence may be caused by phylogenetic rather than trait specific signals. Seven candidate positions of 154 total positions showed changes in CAM species that were restricted to specific phylogenetic groups. One example was position 622 in DPE2, where valine was conserved across 83% of the C_3_ species but changed to leucine in half of the CAM species. This change was detected for DPE2 in all monocotyledonous CAM species, but in none of the eudicotyledonous CAM species. In the C_3_ species, leucine also replaced valine at 622 for three species, all of which belong to monocotyledonous C_3_ species, indicating that leucine at this position is most probably a monocot-specific change that is not functionally relevant in terms of the photosynthetic type (supplemental dataset 11). The percentage of changed sequences in the CAM species and subgroups in which these changes occurred were not the same for all detected changes within one protein. For example, DPE2 was found to have twelve fuzzily conserved changes. While all Orchidaceae shared a change from aspartic acid to nonpolar AAs at position 1311, only two of seven Orchidaceae CAM species possessed a change at position 1558 from lysine to arginine.

### Post-translational modifications

To address the second hypothesis regarding the potential of post-translational modifications being involved in the regulation of CAM we analyzed *in vivo* phosphorylation sites in *K. laxiflora*. Using the same plant material of *K. laxiflora* as for protein abundance analyses, phospho-enrichment and MS analysis were performed and resulted in the identification of 1,805 phosphorylated peptides of which 911 were quantified in at least 3/5 replicates of one time point. At zt8, 647 phosphorylated peptides were quantified in at least 3/5 replicates and at zt20, 805 phosphorylated peptides were quantified in at least 3/5 replicates. Considering only phosphopeptides that were detected at both timepoints and testing for differential abundance of the phosphorylated peptides between light and dark resulted in nine significantly differentially abundant phosphorylated peptides with a q-value<0.05. Eighteen phosphorylated peptides were exclusively detected at zt8 with missing values in all dark samples, and 99 phosphorylated peptides were only detected at zt20 with missing values in all zt8 samples.

Thirty phosphorylated peptides belonging to CAM proteins were identified in at least one replicate of *K. laxiflora* leaf samples after phospho-enrichment and LC-MS/MS analyses. Sixteen of them were identified in at least three of five replicates at zt8 or zt20 and could be tested statistically (= after filtering; supplementary table 9). Four phosphorylated peptides belonging to two isoforms of *Kl*β-CA3 and four phosphorylated peptides belonging to three isoforms of *Kl*GAPC were detected. One phosphorylated peptide of PGM was detected, and it contained two phosphorylated residues, namely Thr-21 and Ser-22, and one peptide matching two *Kl*PPC isoforms with a phosphorylation site at Ser-8 was detected. For *Kl*RP1, two phosphorylated peptides with phosphorylations at positions 75 and 95 were detected. For Starch Excess 4 (SEX4), three phosphorylated peptides were detected with phosphorylations at positions 57, 58, and 64. One phosphorylated peptide matching Phosphoglucan, Water Dikinase (PWD, position 861) was detected (suppl. Something, Figure 5). Analysis with LIMMA showed that Ser-8 in *Kl*PPC was differentially phosphorylated between dark and light, with higher abundance of the phosphorylated peptide at zt20 (adjusted p-value 0.06, log_2_FC 1.33). Additionally, the phosphorylation site at position 71 in RP1 was only detected in the dark samples and missing in all light period replicates (supplementary table 9).

**Figure 5:**
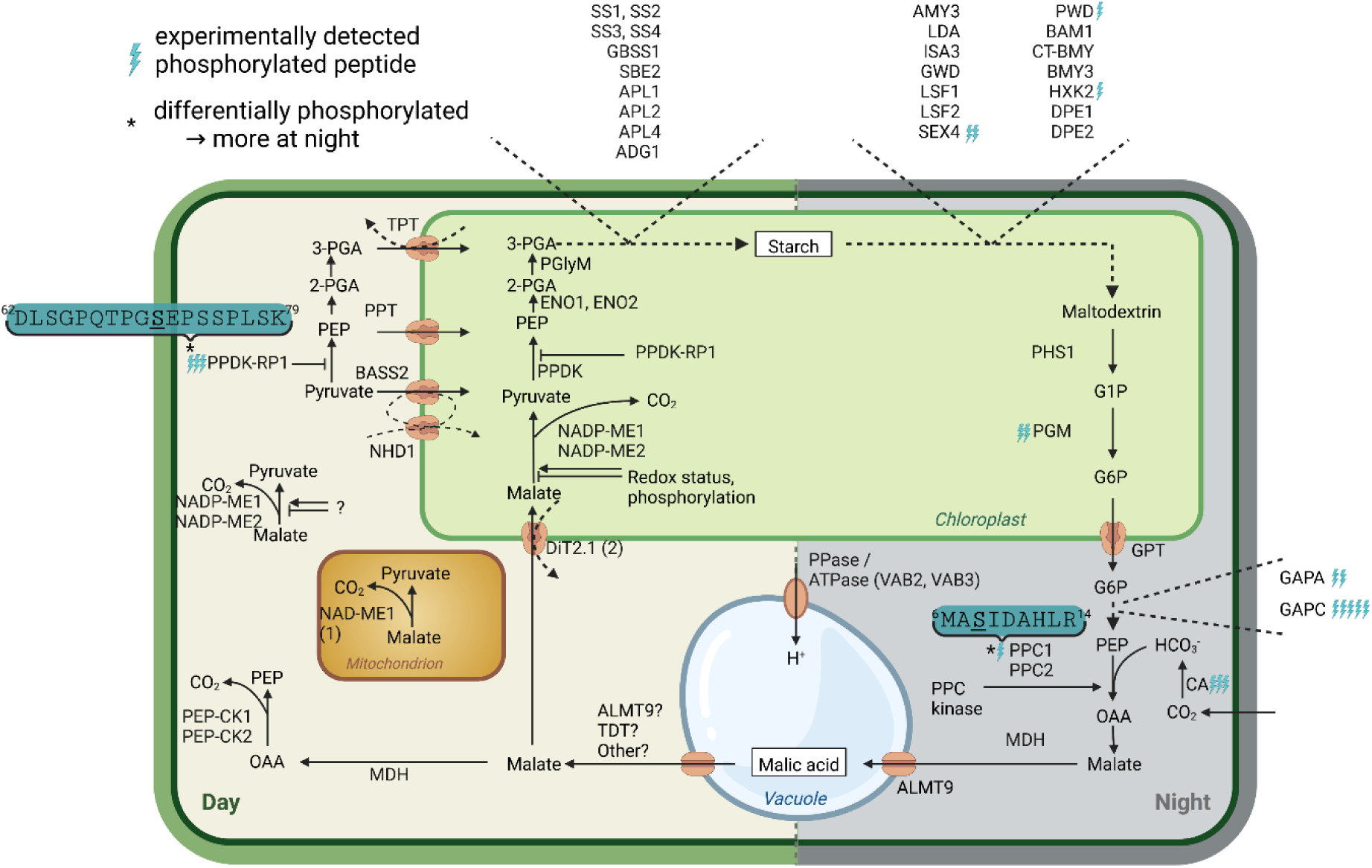
In vivo phosphorylation sites in K. laxiflora. Teal arrows indicate phosphorylated peptides that were identified by MS-analysis of K. laxiflora protein extract sampled at zt8 or zt20. Asterisks mark significantly differential phosphorylation between light- and dark-samples with q-value < 0.05. Significantly differentially abundant phosphopeptides are shown with phosphorylated amino acids underlined. Abbreviations see figure 4.

We analyzed whether the differentially phosphorylated sites were conserved across 47 angiosperm species and found that Ser-8 in *Kl*PPC1 is indeed present in 76% of the orthologous sequences. Most orthologues (86%) of RP1 do not share the Ser-71 site indicating that this might be a *Kalanchoë*-specific phosphorylation site also shared by *K. fedtschenkoi* RP1. In total, five of the 30 detected phosphorylation sites in CAM proteins in *K. laxiflora* were also conserved across 47 angiosperm species (defined as present ≥70%) and were also detected as phosphorylated in *A. thaliana* (database: FAT-PTM). Besides Ser-8 in *Kl*PPC, Ser-124 in *Kl*PGM3, Ser-212 and Thr-213 in *Kl*GAPC and Thr-275 in *Kl*GAPA-2 were conserved and annotated as phosphorylated in their respective *A. thaliana* orthologues (Cruz et al., 2019). Three more sites were conserved, but not annotated as phosphorylated in *A. thaliana*, comprising Ser-58 in *Kl*SEX4, Thr-21 and Ser-22 in *Kl*PGM3. These novel phosphoproteome patterns identified in *K. laxiflora* indicate that phosphorylation beyond the known switch at Ser-8 in PPC may play important roles in the regulation of key enzymes with the CAM pathway.

To test the potential of other protein modifications and the potential for additional conserved phosphorylation sites for CAM species other than *K. laxiflora*, the conservation of phosphorylation, acetylation and S-nitrosylation sites known from *A. thaliana* was tested among CAM proteins. From the 78 proteins that could function in CAM, 63 *A. thaliana* orthologues had at least one phosphorylation, acetylation, or S-nitrosylation PTM site annotated in the database FAT-PTM. In total, 403 PTM sites comprising 180 acetylation sites, 196 phosphorylation sites, and 27 S-nitrosylation sites were analyzed. For these sites, we determined whether the respective AA was conserved in an alignment of the orthologues from 47 angiosperm species (Figure 6a). About half (192/403) of the AAs at PTM sites were conserved (in at least 70% of the orthologous sequences). Conservation of the AA was highest for S-nitrosylation with ∼78% (21/27) of the positions conserved and second highest for acetylated lysines with ∼58% (104/180) of the positions being conserved (Figure 6b). Phosphorylation sites were the most frequent annotated PTM in the analyzed *A. thaliana* proteins, less than a third (64/196) had an AA conservation in 70% or more of the species orthologues in the alignment (Figure 6b). In comparison to phosphorylation sites, acetylation sites (lysines) tend to be more conserved across the 47 species analyzed (Figure 6b).

**Figure 6:**
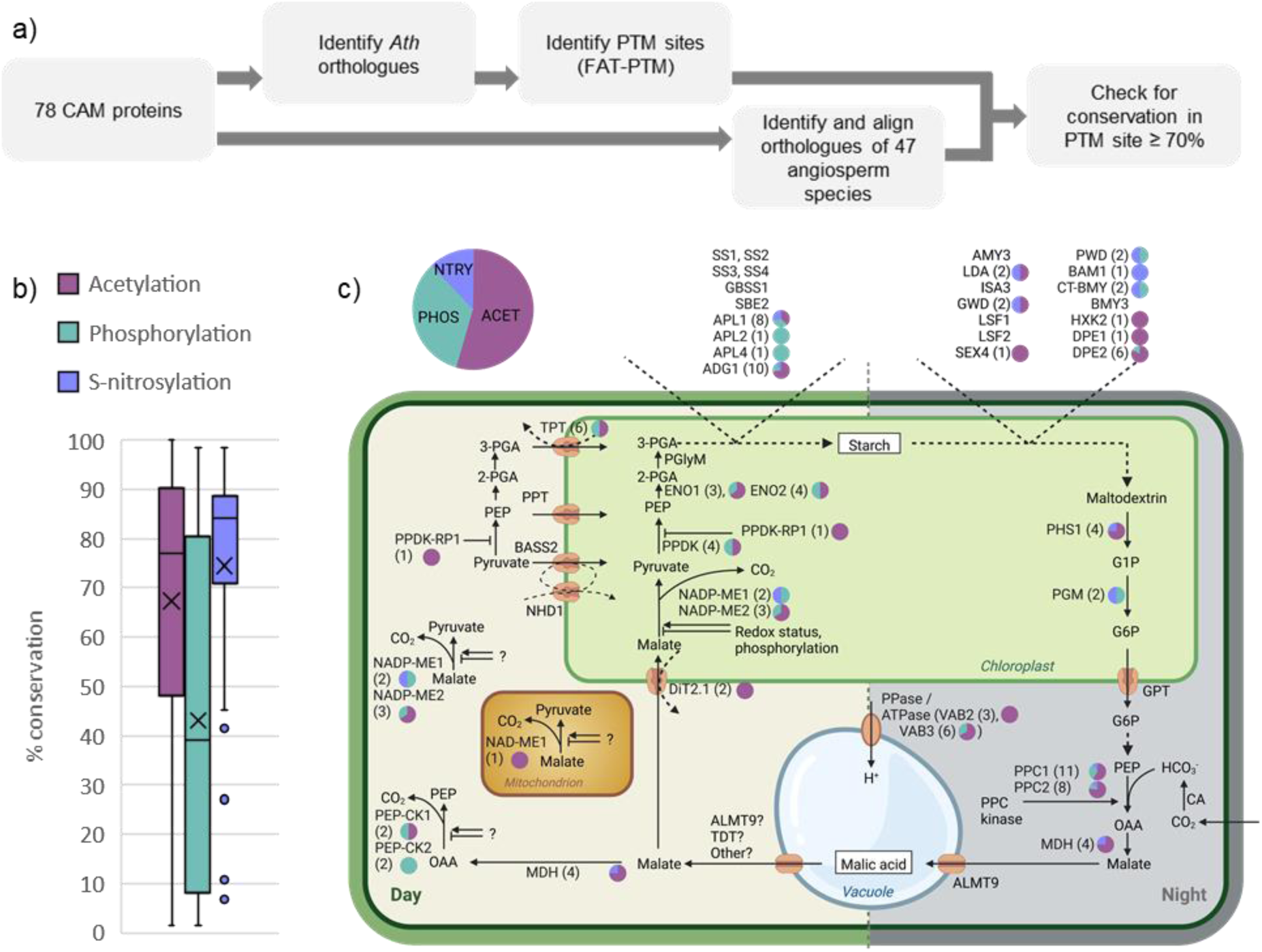
Putative post-translational modification sites. (a) Workflow (b) Boxplot showing the distribution of the percentage of conservation of AAs that are targets to post-translational modification (PTM) in Arabidopsis according to FAT-PTM (Cruz et al., 2019). Orthologues of 47 species were aligned and the conservation of AAs was determined. In total, 403 PTM sites (180 acetylation sites (burgundy), 196 phosphorylation sites (teal), and 27 S-nitrosylation sites (purple)) were analyzed from 63 Arabidopsis proteins. (c) CAM pathway with conserved putative PTM sites. Number in brackets gives the total amount of acetylations, phosphorylations, and S-nitrosylations with the amino acid (AA) being conserved in at least 70% across 47 angiosperm species. Little pie charts indicate the proportion of acetylations (ACET, burgundy), phosphorylations (PHOS, teal), and S-nitrosylations (NTRY, purple). Pie chart on the top left indicates the proportions of conserved putative PTM sites in all analyzed CAM proteins. Abbreviations: see figure 4.

The conserved putative acetylation, S-nitrosylation and phosphorylation sites were identified throughout the whole pathway (Figure 6c). Of all proteins, 33 had no conserved putative acetylation, S-nitrosylation, or phosphorylation site. Putative acetylation sites with the lysine conserved across 47 angiosperm species were found in 31 proteins including proteins involved in carboxylation, vacuolar malate storage, decarboxylation and the starch module. Conserved positions for S-nitrosylation were detected in 18 proteins. Nine of these proteins had conserved sites for both, acetylation and S-nitrosylation (Figure 6c and supplemental Table 9).

## Discussion

### Transcripts and Proteins of CAM genes do not show on/off patterns

CAM has the potential to be regulated at several different molecular levels. To test the influence of diurnal abundance patterns of CAM transcripts and proteins, their abundance was quantified in the middle of the dark and light periods in the obligate CAM plant *K. laxiflora*. As expected from diurnal time courses, 6,247 transcripts and 234 proteins were detected as being differentially abundant between light (zt8) and dark (zt20) (Figure 1b, c). GO term analysis showed typical patterning of differentially abundant transcripts, for example, with the GO term photosynthesis being less abundant at night (e.g. Heyduk et al., 2019; supplemental Table 6). The transcriptome analysis revealed that 42 out of 84 candidate CAM-associated transcripts were differentially abundant between light and dark (Figure 2a), consistent with the results of time-course transcriptome data in other CAM species (Ming et al., 2015; Abraham et al., 2016; Heyduk et al., 2019; Wai et al., 2019). Within the carboxylation/decarboxylation cycle one *PPC* isoform was highly expressed at ∼100 tpm during the night (Figure 2a). For decarboxylation, one *NAD-ME1* and two *NAD-ME2* isoforms were highly expressed at 395 and 122-130 tpm during the day. Since the most abundant *NADP-ME* isoform was measured at 100 tpm and the most abundant PEP-CK isoform was at 6 tpm during the light, the data suggests that decarboxylation mostly occurs via NAD-ME possibly with contributions from the other decarboxylation enzymes. This finding fits very well with previous work using stable transgenic mutant *NAD-ME* RNAi lines of very closely related species *K. fedtschenkoi*, which demonstrated clearly that mitochondrial NAD-ME was essential for a functioning CAM cycle in *Kalanchoë* (Dever et al., 2015). Importantly, NADP-ME activity was not able to rescue the inability of the *NAD-ME* RNAi lines to decarboxylate malate in the light (Dever et al., 2015). In *T. fruticosum*, the primary decarboxylation route could not be predicted from the transcript abundance data (Brilhaus et al., 2016). In C_4_ plants, a major ME can usually be proposed based on transcript abundance, sometimes with contributions by PEP-CK (Bräutigam et al., 2014). Unlike in C_4_, in CAM, no diffusion gradient between cell types is required, which may enable the contribution of two different malic enzymes. When compared between zt8 and zt20, *PPC1* transcript abundance was significantly different in *K. laxiflora*, but transcripts at zt8 remained at 50 tpm. A similar pattern has been reported previously in other CAM transcriptomic analyses, for example in *Ananas comosus, Agave americana, K. fedtschenkoi*, and *S. album*, or *Tillandsia* species (Ming et al., 2015; Abraham et al., 2016; Yang et al., 2017; Wai et al., 2019; Groot Crego et al., 2024). The authors show that transcript abundance changes throughout the diurnal cycle but never represents a binary on/off pattern for the CAM enzymes. Protein abundance was found here to be stable for all PPC isoforms, and the protein was present during the day (Figure 2b). In *A. americana*, it was shown that PPC protein abundance peaked with maximal activity, but the protein was present throughout light and dark periods (Abraham et al., 2016). Similar results were found for *K. fedtschenkoi* mesophyll cells where PPC protein abundance peaked at night but protein was also present during the day (Abraham et al., 2020). The presence of both transcript and protein at a time point where flux through the CAM pathway is off suggests that an additional regulation layer is required on top of the small (Abraham et al., 2016; Abraham et al., 2020), if any (Figure 1b) PPC protein abundance changes. Peak transcript abundance of the other metabolic enzymes in *K. laxiflora* for the major isoform matched peak activity but again, substantial transcript amounts remained during at the opposite time when activity is predicted to reach its 24-h minimum (Figure 2). Similar observations have been reported for other independent origins of CAM including *A. americana*, (Abraham et al., 2016), *A. comosus*, (Sharma et al., 2017), and *S. album* (Wai et al., 2019). The timing of peak transcript abundance frequently matched the timing of peak activity, but transcript abundance was never near zero during off-times. A highly resolved time-course transcriptome set from the closely related species *K. fedtschenkoi* (JGI data release, project ID 1155736) underlined these conclusions (supplemental figure 3). The transcripts encoding core CAM pathway proteins had stable abundance when compared between the middle of the light and dark (Figure 2b), which is entirely consistent with previously reported findings for *Agave* and *Kalanchoë* (Abraham et al., 2016; Abraham et al., 2020). The only CAM protein that was significantly differentially abundant on the transcript level and also detected and differentially abundant on the protein level in this study was the malate/ fumarate transporter (TDT). Strong up-regulation of this tonoplast-localized organic acid transporter at in the light period sample supports the previously proposed hypothesis that this transporter might be involved in the export of malate from the vacuole for decarboxylation in *K. laxiflora* (Borland et al., 2009).

When also considering the CAM genes *sensu lato*, that is, including the enzymes of carbon storage, 20 of the 84 transcripts were strongly significantly differentially abundant between light and dark, with |log_2_FC|>1 (Figure 2a). In eleven of these cases, the differential abundance matched the expected time of activity. For example, *GBSS1* was significantly up-regulated on the transcript level during the light, at a time point when enzyme activity is also predicted to peak as part of the pathway for light-period starch synthesis (Streb and Zeeman, 2012). Further examples of such correlations included *APL4*, *SBE2*, and *ADG1,* which all encode enzymes within the starch synthesis pathway (Streb and Zeeman, 2012), and all displayed differential transcript abundance that was higher in the light (Figure 2a). In contrast, seven potential CAM-associated transcripts were found here to have higher abundance at the time point when lower activity of the encoded enzyme is predicted. For example, *GPT* encodes a glucose 6-phosphate (G6P): phosphate translocator that localizes to the chloroplast envelope. G6P is predicted to be the main sugar exported to the cytosol in the dark period during phosphorolytic starch breakdown, and this translocator may function in either the dark, for G6P export from the chloroplast, and/ or in the light for G6P import during pyruvate recycling to starch (Borland et al., 2009; Ceusters et al., 2021). However, here transcripts for one *GPT* gene were significantly upregulated at zt8, with a log_2_F change of 1.4 (Figure 2). This isoform corresponds to *GPT2* in *A. thaliana*, which has been shown to be upregulated under high light stress to enhance export of sugar phosphates (Boxall et al., 2020; Balcke et al., 2023). Peptides for all CAM pathway enzymes *sensu lato* were also detected for both timepoints except for SS3, where true binary protein presence/absence was detected (Figure 2b).

Among all putatively CAM-associated transcripts analyzed, there was one notable exception to the rule that transcripts were detected in both the light and dark sample. Transcripts from two *PPCK* genes, *PPCK1* and *PPCK2*, were highly upregulated at zt20 (log_2_F changes of 9.9 and 5.3, respectively), with zt8 transcript levels at or below 1 tpm (Figure 2a). Transcripts for a third *PPCK* gene, *PPCK3*, displayed the opposite pattern of differential abundance, with a four-fold up-regulation in the light sample relative to the dark (Figure 2a). *PPCK1* in *Kalanchoë* has been demonstrated using RNAi mutant transgenic lines to be essential for PPC phosphorylation in the dark period, and in its absence nocturnal CO_2_ assimilation was reduced to 33% of the wild type level (Boxall et al., 2017). Whilst *PPCK1* and *PPCK2* were detected here as being more abundant in the dark, consistent with a function in regulating CAM PPC, *PPCK2* was not required for PPC phosphorylation in *K. fedtschenkoi* as the PPCK1 RNAi transgenic lines had not detectable phospho-PPC (Boxall et al., 2017). Thus, the function of *PPCK2* in *Kalanchoë* leaves remains to be elucidated, with one intriguing possibility being that it works to regulate PPC in other cell types of the leaf, not in the mesophyll cells where the reactions of the CAM cycle occur. Likewise, *PPCK3* that peaked in the light period may work in other cell types or may regulate the house-keeping PPC isoform that functions in the replenishment of 4-carbon skeletons used for amino acid synthesis. Consistent with this, *PPCK* transcripts were induced by light in the C_3_ plants barley and *A. thaliana* (Hartwell et al., 1996; Fontaine et al., 2002), and absence of PPC phosphorylation in *A. thaliana* leaves had pleiotropic effects on the growth of the plant (Meimoun et al., 2009). Similar data, with a large increase in *PPCK1* transcript abundance in the dark period, was also reported for other CAM transcriptomic studies. For example, *PPCK1* transcripts peaked in the dark in data for *K. fedtschenkoi* (Yang et al., 2017; JGI data release) and *S. album* (Wai et al., 2019). However, no peptides for PPCK were detected in the MS data (Figure 2b), which is frequently the case for regulatory proteins of low abundance (Bantscheff et al., 2007). Furthermore, the failure to detect PPCK peptides within the MS data for proteins extracted from either light or dark leaves is consistent with earlier estimates of PPCK protein abundance in CAM leaves of *Kalanchoë* from attempts to purify PPCK to homogeneity from CAM leaves (Hartwell et al., 1999). PPCK purification studies estimated that, even at the time of peak activity, PPCK protein represents less than 1 in 10^5^ of the soluble protein of the leaves of *K. fedtschenkoi* (Hartwell et al., 1999).

The quantitative results presented here for transcript and peptide abundance changes suggest that, in *K. laxiflora*, other molecular mechanisms must operate to optimize metabolic flux through the CAM cycle relative to the light/dark cycle (Figure 2). This finding contrasts with data for C_4_ species, where transcript abundance changes have been found to be diagnostic for identifying C_4_ genes (Bräutigam et al., 2011; Christin et al., 2012), and spatial separation of both transcripts and proteins between mesophyll and bundle sheath cells was detectable (Majeran et al., 2005; John et al., 2014). Schiller and Bräutigam (2021) proposed that CAM is a continuous rather than a binary trait and that regulation of CAM is likely only partially caused by circadian changes of abundance of CAM transcripts and proteins thereby implicating that complex/ extensive rewiring of the circadian clock is likely not necessary in CAM plants. In response, Winter and Smith (2021) showed that detectable malic acid accumulation and therefore CO_2_ fixation during the night is a feature exclusive to CAM plants. They conclude that because of tight connection of the circadian clock and dark CO_2_ fixation rewiring of the circadian clock to metabolic processes would be indeed necessary in CAM plants. Our analyses in *K. laxiflora* demonstrate that, similar to studies in other CAM species (Ming et al., 2015; Abraham et al., 2016; Yang et al., 2017; Wai et al., 2019), diurnal variation in metabolic CAM genes is detectable (Figure 2) but not binary, that is, not in an on/off pattern. In contrast, the oscillation of the regulatory PPCK is binary and observed transcript abundance changes were sufficient to explain activity patterns in CAM, with activity of the kinase at night leading to enhanced PPC phosphorylation, CO_2_ fixation and malic acid accumulation (Hartwell et al., 1999; Boxall et al., 2017). The data suggests that transcriptional regulation of metabolic CAM genes by circadian clock or diurnal regulators might be necessary (Ming et al., 2015; Abraham et al., 2016; Yang et al., 2017; Wai et al., 2019, Figure 2a), but is certainly not sufficient for regulation (Figure 2a, b). Circadian or diurnal regulation of regulatory genes such as PPCK, due to their comparatively lower protein abundance (Figure 2b), are suitable to direct flux as proposed earlier (Hartwell et al., 1999; Boxall et al., 2017; Schiller and Bräutigam, 2021).

### CAM plants have accumulated conserved putative adaptive changes in the starch module

CAM plants need to accommodate nightly storage of malate and daily storage of the sugars from photosynthesis and pyruvate/ PEP recycling, as well as higher fluxes towards these storage metabolites and hence through the CAM cycle. We hypothesized that adaptive mutations may have accumulated in CAM proteins. Since CAM represents a continuum of CAM types, including the recruitment of alternative pathways for malate decarboxylation and starch/ soluble sugar storage, depending on the species (Silvera et al., 2010), we further hypothesized that conserved changes and adaptive mutations in AA sequences of CAM proteins would only be detectable fuzzily (i.e. not always present) across a diverse taxonomic sampling of independent CAM origins. Data for obligate CAM species such as *K. fedtschenkoi* or *K. laxiflora,* and for inducible CAM cycling species such as *S. album,* in which CAM-cycling is induced by drought stress (Wai et al., 2019), or *T. fruticosum*, in which CAM is induced by drought stress (Brilhaus et al., 2016) were included. The degree of CAM and the inducibility can only be determined via malic acid accumulation and direct measurement of nocturnal CO_2_ assimilation and stomatal opening (Winter, 2019), which requires specialized equipment as CAM plant leaves are usually fleshy (Figure 3a-e) and do not fit standard gas exchange equipment. Hence, the degree and inducibility of CAM are not known for all species under investigation. CAM species also use different biochemical routes in the CAM pathway. Two main routes are described for decarboxylation of organic acids, either ME and PPDK, or MDH and PEP-CK (Gilman and Edwards, 2020), or a mix of several enzymes (Christopher and Holtum, 1996). *K. fedtschenkoi* uses mitochondrial NAD-ME and PPDK (Dever et al., 2015; Hartwell et al., 2016; Yang et al., 2017), and this was supported here in the transcriptomic data for *K. laxiflora* (Figure 2a). In contrast, *A. comosus* uses MDH and PEP-CK for malate decarboxylation in the light, and this was supported by quantitative transcriptome data from a diel timecourse of leaf samples (Ming et al., 2015). CAM species also differ in their main transitory carbohydrate pool for provision of the PEP required as the substrate for CO_2_ assimilation by PPC in the dark. *K. fedtschenkoi* stores mainly starch in the chloroplast, and *A. comosus* mainly soluble sugars in the vacuole (Ming et al., 2015; Yang et al., 2017; but see Antony et al., 2008). Information on the main carbohydrate pool is often derived from physiological analyses like measuring starch and soluble sugar content at regular intervals over the light and dark period, and/ or during the progression of CAM induction by abiotic stress, especially drought, as for example reported for *S. album* (Wai et al., 2019). Experimental data that would allow the subtype of CAM (obligate/ inducible, decarboxylation route, carbohydrate pool) to be categorized reliably is lacking for many species, and since enzymatic assays for the possible decarboxylation enzymes NAD-ME, NADP-ME and PEP-CK suggest that species exist that use more than one decarboxylation enzyme simultaneously, grouping of CAM species into different categories remains inaccurate (Christopher and Holtum, 1996). Here, CAM species spanning all of the mentioned sub-types were included, but fuzziness was allowed, with only 50% conservation of a specific AA residue required in the detection of adaptive mutations in the hope that specific adaptations that may be unique to only some sub-types of CAM would not be missed. Using 20 CAM species, 154 AA residue positions in 31 CAM proteins were found to have a fuzzy change in their AA sequence in comparison to the orthologous sequence in C_3_ species, but an uneven distribution around the 24-h CAM cycle biochemical steps was observed (Figure 4d). Fuzzily conserved changes were observed mostly in enzymes that function in starch synthesis and degradation, and not in the CAM carboxylation and decarboxylation steps of the pathway (Figure 4d).

Goolsby et al. (2018) analyzed species in the portullugo clade (Caryophyllales) for AA changes/ adaptive mutations between C_3_ and obligate or inducible CAM species. They focused on CAM genes *sensu stricto* and did not find changes between the AA sequences of CAM genes in C_3_ and CAM species. Likewise, here, even with an expanded and far greater diversity of species that spanned diverse monocot and eudicot orders, no conserved AA changes in the CAM genes *sensu stricto* were discovered, apart from one exception in PEP-CK where a serine to cysteine change was conserved in eleven CAM species (supplemental dataset 21). However, due to the lack of definitive information categorizing the decarboxylation pathway biochemical sub-types of CAM in each species, it was not possible to determine whether this change occurs only in CAM species that use MDH and PEP-CK for malate decarboxylation in the light. The PEP-CK AA change was detected in the orthologue in *T. fruticosum* where transcriptomic data suggested ME was the enzyme used for decarboxylation of malate following the induction of CAM by drought (Brilhaus et al., 2016). All other analyzed CAM genes *sensu stricto* did not yield any fuzzily conserved AA changes between C_3_ and CAM species, indicating that these genes are regulated on other levels or at least have not needed to evolve changes in their AA sequence and therefore structure for their CAM-specific function in many CAM species. Yang et al. (2017) changed single AAs in PPC2 of *K. fedtschenkoi* and the orchid *Phalaenopsis equestris,* and discovered this changed the properties of the enzyme when it was heterolgously expressed and assayed, which supported the conclusion that, in single species, single changes may be adaptive for optimal CAM. However, *PPC1* is the only *PPC* gene family member necessary for CAM in *Kalanchoë*, and in the absence of *PPC1*, *PPC2* did not complement the mutation and rescue the CAM phenotype, despite *PPC2* transcripts being induced in the *PPC1* RNAi lines (Boxall et al., 2020). The most strongly silenced PPC1 RNAi line lacked detectable PPC activity in mature CAM leaves and the plants performed no detectable nocturnal atmospheric CO_2_ assimilation (Boxall et al., 2020). Such functional studies using transgenic mutants emphasize the crucial need for systematic mutant studies of candidate CAM-recruited members of each CAM gene family in order to confirm AA sequence level changes that may be identified across multiple species are linked with the CAM-recruited member of each CAM gene family.

Here, the breadth of CAM species sampling was extended to 7/15 orders with known CAM origins, spanning both the Monocots and Eudicots (Figure 3g). The broad sampling together with the fuzziness criterium inevitably led to phylogenetic bias to some degree. Orchidaceae and Cactaceae were over-represented within the CAM species (Figure 3g). In seven cases, phylogenetic bias was detected, namely in the proteins DPE2, AMY3, SS1, SS3, and TDT (supplemental dataset 4, 11, 29, 30, 32), which highlights that phylogenetic bias was only a minor contributor to the conserved AA changes detected here (Figure 4d).

When taking CAM genes *sensu lato* into account, which expanded the set of analyzed genes to include starch synthesis and breakdown modules, as described in Schiller and Bräutigam (2021), the accumulation of putatively adaptive and conserved changes was detected in the group of proteins involved in starch synthesis and breakdown (Figure 4d). Feike et al. (2021) reported that a single AA mutation in β-amylase caused an accelerated starch degradation phenotype *in vivo*, possibly by higher binding affinity for its interaction partner, leading to more efficient starch degradation. In the analysis presented here this AA change was not detected, but several other conserved changes in putative CAM copies of proteins involved in starch synthesis and breakdown were detected. Two thirds of the proteins involved in carbohydrate storage and breakdown (21/30) had fuzzily conserved changes in their AA sequence between CAM and C_3_ species. Among these proteins were PHS1 and DPE. PHS1 is a key player in phosphorolytic starch degradation. It catalyzes the phosphorolysis of glucose residues to yield glucose-1-phosphate (Smith and Zeeman, 2020). Upon downregulation of the chloroplastidic PHS1 orthologue in stable transgenic RNAi lines of the obligate CAM plant, *K. fedtschenkoi*, the transgenic lines re-routed starch degradation via the amylolytic route and had reduced dark CO_2_ uptake, reduced water-use efficiency, and an overall reduction in growth (Ceusters et al., 2021). Two AA positions in PHS1 that were conserved in C_3_ species were found to differ in > 50% of the analyzed CAM sequences (Figure 4d, supplement excel). The AA sequence changes shared in CAM species were found in the same subset of species belonging to Orchidaceae and Saxifragales, Crassulaceae. In both positions, a conserved polar AA in C_3_ PHS1 was switched to a non-polar and acidic AA, respectively, in half of the analyzed CAM sequences. Among these species is *K. fedtschenkoi*, which was analyzed in the study presented by Ceusters et al. (2021). The AA positions identified here in PHS1 together with the importance of this enzyme as a possibly rate-limiting step for nocturnal carboxylation in CAM (Borland and Dodd, 2002; Cushman et al., 2008a; Haider et al., 2012), suggest that it may be important to include starch cycle engineering using CAM protein isoforms into efforts to engineer CAM into non-CAM species to enhance their water-use efficiency. The re-routing of the pathway of starch degradation in the dark period in CAM species may have been favored during the evolution of CAM due to the capacity for CO_2_ assimilation by PPC being dependent on nocturnal supply of PEP and the increased energetic demands of CAM compared to C_3_ (Borland et al., 2016; Shameer et al., 2018). Export of G6P from the chloroplast during nocturnal starch degradation offers a specific energetic advantage since glycolytic conversion of G6P to PEP in the cytosol provides ATP (Ceusters et al., 2021). In contrast, the amylolytic route of leaf transitory starch breakdown that is used by C_3_ *A. thaliana* in the dark leads to the export of maltose and glucose from the chloroplast, and this in turn leads to the requirement for glucose phosphorylation by hexokinase in the cytosol, which consumes ATP, before sugar phosphates can be converted to PEP through the rest of glycolysis. Modelling of the CAM pathway using a flux-balance approach suggested that the use of the phosphorolytic over the amylolytic pathway for nocturnal starch breakdown could save 14 – 26% of nocturnal ATP consumption in CAM (Shameer et al., 2018).

DPE1 and DPE2 are also important for starch breakdown. Borland et al. (2016) and Ceusters et al. (2019) have shown that DPE2 activity was low in the CAM plants *M. crystallinum*, *Phalaenopsis* ‘Edessa’ and *K. fedtschenkoi,* which was assumed to be a characteristic of CAM species. DPE2 is involved in hydrolytic starch breakdown by metabolizing exported maltose, but is not involved in phosphorolytic starch breakdown. Here, 12 AA changes were identified in the analyzed DPE2 orthologues (supplemental dataset 11). The detected changes in DPE2 from CAM species *P. equestris* and *K. fedtschenkoi* were not identical, which may be due to the independent origins of CAM in the different families analyzed, and may point to multiple evolutionary routes to the same end. In comparison to that, DPE1, which is involved in both starch breakdown pathways, was more active in *K. fedtschenkoi* and *M. crystallinum* than in *A. thaliana* in terms of the total extractable activity measured *in vitro* (Borland et al., 2016; Ceusters et al., 2019). There were also 12 changes in the AA sequence found for DPE1. Overall, the large number of fuzzily conserved changes in starch synthesis and breakdown suggest that the AA sequence and protein structure of enzymes involved in starch metabolism may have adapted during the evolution of CAM in the majority of CAM species.

In conclusion, 154 AA positions were identified here as having fuzzily conserved changes in CAM proteins when 20 CAM species are compared with 27 C_3_ species with many of the identified AA changes clustering in particular proteins. The high number of up to 21 fuzzily conserved changes per protein and the fuzzy conservation itself also suggests that each modification only contributes a small adaptation to the CAM pathway. In CAM engineering into C_3_ efforts, it will therefore likely be critical to lift and transfer the whole pathway from one species to make sure that a fitting complement of modifications is transferred.

### Phosphorylation and other post-translational modifications have high potential to regulate CAM

The two tested control mechanisms, abundance of transcripts and proteins and changes to the protein sequence, cannot alone serve to control flux through the CAM pathway. The abundance pattern was not binary (Figure 2), and the changes to the AA sequence of CAM proteins clustered in the, mostly chloroplast-localized, starch synthesis and breakdown pathways (Figure 4d). Post-translational modifications provide another avenue to control the pathway, and are in fact the most thoroughly understood mechanisms for the temporal control of the 24-h CAM cycle. For example, characterization of transgenic RNAi lines of *Kalanchoë* lacking detectable levels of PPCK and with no nocturnal PPC phosphorylation displayed a 66% drop in nocturnal CO_2_ assimilation relative to the wild type, and the normally robust and high-amplitude free-running circadian rhythm of CO_2_ assimilation collapsed to arrythmia (Boxall et al., 2017). We therefore analyzed the *in vivo* phosphorylation status of proteins in CAM leaves of *K. laxiflora* using samples collected in the middle of the light and dark period. We detected 1805 non-redundant Phosphopeptides in 806 proteins. This is comparable to the 2175 phosphopeptides in 1255 proteins detected in a study reported for the C_3_ species *A. thaliana* (Nukarinen et al., 2016), and comparable to the 2232 phosphosites detected in 401 proteins in rice (Hamzelou et al., 2021). The analysis here focused on the proteins in the CAM pathway. A well-known PTM site with regulatory function within the CAM pathway is the circadian clock-controlled nocturnal phosphorylation of a conserved N-terminal serine residue in PPC (Boxall et al., 2017). In the phosphoproteomic data collected here, the number of phospho-PPC N-terminal peptides detected displayed differential abundance between the dark and light, with roughly 2.5-fold higher abundance of the phosphorylated peptide at zt20 (supplemental table 9). In *A. thaliana* plant-type PPC, this regulatory phosphorylation site occurs at Ser-11, whereas in *K. laxiflora* CAM PPC1, the phosphorylatable serine is at Ser-8. Aligning PPC orthologues of 47 angiosperm species showed that serine was conserved in ∼74% of the sequences at this position. As predicted by previous work, the phosphorylation of *Kl*PPC1 at Ser-8 was significantly more abundant at zt20 in comparison to zt8, highlighting the known mechanism for regulating the allosteric properties of PPC, with Ser-8 phosphorylation leading specifically to a 10-fold decrease in the sensitivity of the enzyme to feedback inhibition by L-malate (Nimmo et al., 1986; Boxall et al., 2017; Li et al., 2023). Protein abundance of PPC and total extractable specific activity remained remarkably stable across the light/dark cycle in *Kalanchoë* (Figure 2b; Nimmo et al., 1984).

The second phosphorylation site detected as being differentially abundant between the light and the dark was in the PPDK regulatory protein, RP1, which can function both as a PPDK kinase and PPDK phosphatase. This RP1 phosphosite was not shared between the 47 angiosperm species, but it was shared with a *K. fedtschenkoi* RP1 orthologue. RP1 might therefore harbor a *Kalanchoë*-specific phosphorylation site, which could allow the regulation of RP1 activity via phosphorylation. The reaction mediated by PPDK is an energetically costly reaction from pyruvate to PEP because the reaction consumes energy in the form of ATP (Chastain et al., 2011). Tight regulation of this step is therefore necessary and this is achieved through the activity of RP1 through its phosphorylation/dephosphorylation of PPDK (Dever et al., 2015). Unlike PPCK, RP1 protein was detected at both zt8 and zt20, but it was not significantly differentially abundant (Figure 2). Phosphorylation of the protein might activate RP1 at night, thereby enabling the phosphorylation of PPDK, which in turn inhibits this reaction in the dark. Given that PPDK-RPs can catalyze both the phosphorylation and dephosphorylation of PPDK, it is tempting to propose based on our findings that the dephosphorylation and activation of PPDK in the light in *Kalanchoë* may be achieved due to the dephosphorylation of RP1; i.e. dephospho-RP1 may function preferably as the PPDK phosphatase, whereas phospho-RP1 may function as the PPDK kinase.

When looking at the conservation of putative phosphorylation sites across CAM proteins overall, proteins localized to both the cytosol and chloroplast were identified as having conserved AAs that might be phosphorylated (Figure 6c). Phosphorylation is mediated by protein kinases and regulates enzyme activity, protein stability, protein sub-cellular localization, or protein-protein interactions (Budde and Chollet, 1988). Zhu et al. (2018) have analyzed the kinome of pineapple and found fourteen kinases that are potential regulators involved in photosynthesis or CAM regulation based on their spatiotemporal expression patterns. We analyzed public circadian data of *K. fedtschenkoi* from the same genus as *K. laxiflora* (JGI data release, Project ID: 1155736). A search for the conserved protein kinase domains resulted in the identification of 1,109 candidate protein kinases, and 904 of these had rhythmic, circadian transcript abundance patterns (supplemental figure PCA and Heatmap of kinases). Only one protein kinase with rhythmic transcript abundance pattern was predicted to be plastid localized indicating that proteins in the plastid are perhaps less likely to be regulated by phosphorylation mediated by nuclear-encoded protein kinases that are regulated at the level of the transcript abundance. Other rhythmic protein kinase candidates were mostly predicted to encode cytosolic proteins (supplemental figure PCA and Heatmap of kinases). While the number of rhythmic kinases was too high to allow a candidate test approach, the high number demonstrates the extent of potential kinase-based regulation for metabolic pathways such as CAM.

While phosphorylations have been studied for decades, more recently, additional PTMs like acetylation and S-nitrosylation have been analyzed for their regulatory importance (Finkemeier et al., 2011; Hess and Stamler, 2012). It has been shown that evolutionarily conserved lysines serve as hotspots for acetylation (Balparda et al., 2022). In the *A. thaliana* orthologues of CAM proteins, 180 lysine acetylation sites are annotated in FAT-PTM (Cruz et al., 2019). More than half of these lysines are conserved in the 47 angiosperm species analyzed, including the CAM species (Figure 6b). For example, a putative acetylation site was conserved in the orthologues of PEP-CK, and it has been shown in mice that PEP-CK toggles between carboxylation and decarboxylation depending on its acetylation state (Latorre-Muro et al., 2018). Proteomics analysis of acetylated peptides in a CAM species like *K. laxiflora* might provide insights into whether these PTMs occur in a time-structured manner in CAM proteins over each light/dark cycle, thereby offering the potential for diel and possibly circadian regulation of activity to support optimized timing of flux through CAM.

## Conclusion

The results presented here indicate that CAM regulation happens on multiple levels and is not limited to circadian or diel regulation by *de novo* transcription. Transcript and protein abundances were not found to change in an on/off pattern between light and dark except for the key circadian clock-controlled regulatory protein kinase PPCK. This further cements the pivotal role of this protein kinase in ensuring optimal temporal control of CAM over each 24-h light/dark cycle (Boxall et al., 2017). In CAM species, starch synthesis and degradation enzymes were found frequently to carry fuzzily conserved AA sequence changes that may each contribute a small adaptation to the altered metabolic fluxes that support efficient CAM, but the carboxylation/decarboxylation enzymes lacked conservation of AA residue changes. Post-translational protein phosphorylation occurs *in vivo* in *K. laxiflora* CAM proteins including two significantly differentially abundant phosphorylation sites, of which one is reported for the first time (RP1). Bioinformatic analyses demonstrate the potential for additional modifications such as cysteine nitrosylation and lysine acetylation of conserved AAs, which highlights an area that is ripe for further experimental investigation in the light of the potential that these modifications may be involved in regulation and fine tuning of CAM.

## Additional Information

### Author Contributions

KS produced, analyzed and interpreted data, co-wrote the manuscript, SJ analyzed data, SZ analyzed data, PV produced data, JH provided material, interpreted data, and co-wrote the manuscript, JE produced and analyzed data, IF provided materials, aided in experimental design, supervised, and interpreted data, AB conceived of the study, supervised, interpreted data, and edited the manuscript

## Acknowledgements

The authors gratefully acknowledge Christine Schlüter, Paulina Heinkow, and Marcus Höpfner for expert technical assistance and the CeBiTec compute cluster for computational resources. This research was funded by Bielefeld University and the DFG (instrumentation INST 211/744-1 FUGG). Pathway figures (4d, 5, 6c) were created with BioRender.com.

## Data Availability

RNA-seq reads generated in this study were deposited in the NCBI SRA under BioProject ID PRJNA1172678.

Mass spectrometry raw data and MQ results files, as well as the custom protein FASTA file are available via JPost identifier JPST003223.

